# PP1 and PP2A use opposite phospho-dependencies to control distinct processes at the kinetochore

**DOI:** 10.1101/610808

**Authors:** Richard J Smith, Marilia H Cordeiro, Norman E Davey, Giulia Vallardi, Andrea Ciliberto, Fridolin Gross, Adrian T Saurin

**Affiliations:** Division of Cellular Medicine, School of Medicine, University of Dundee. DD1 9SY; Conway Institute of Biomolecular & Biomedical Research, University College Dublin, Belfield, Dublin 4, Ireland; Division of Cancer Biology, The Institute of Cancer Research, 237 Fulham Road, London SW3 6JB, UK; Istituto Firc di Oncologia Molecolare, IFOM, Milano, Italy

## Abstract

PP1 and PP2A-B56 are major serine/threonine phosphatase families that achieve specificity by colocalising with substrates. At the kinetochore, however, both phosphatases localise to an almost identical molecular space and yet they still manage to regulate unique pathways and processes. By switching or modulating the positions of PP1/PP2A-B56 at kinetochores, we show that their unique downstream effects are not due to either the identity of the phosphatase or its precise location. Instead, these phosphatases signal differently because their kinetochore recruitment can be either inhibited (PP1) or enhanced (PP2A) by phosphorylation inputs. Mathematical modelling explains how these inverse phospho-dependencies elicit unique forms of cross-regulation and feedback, which allows otherwise indistinguishable phosphatases to produce distinct network behaviours and control different mitotic processes. Therefore, the kinetochore uses PP1 and PP2A-B56 because their binding motifs respond to kinase inputs in opposite ways. Genome-wide motif analysis suggests that many other pathways also select for these same key features, implying that these similar catalytic enzymes may have diverged during evolution to allow opposite modes of phospho-regulation.

## Introduction

Protein phosphatase 1 (PP1) and protein phosphatase 2A (PP2A) are large phosphatase families that are responsible for most of the serine/threonine dephosphorylation in eukaryotic cells (Brautigan and Shenolikar, 2018; Heroes et al., 2013). This is exemplified by the fact that PP2A inhibition causes approximately half of the phosphorylation sites in the human proteome to change significantly (Kauko et al., 2018). PP1 and PP2A use structurally-related catalytic domains that are remarkably well-conserved and relatively promiscuous in vitro (Ingebritsen and Cohen, 1983). In vivo, however, they are believed to achieve specificity by interacting with short linear motifs (SLiMs) that localise them to their required sites of action (Brautigan and Shenolikar, 2018). The best characterised SLiM for PP1 is the RVxF motif that is present in approximately 90% of the validated PP1 interacting proteins (Heroes et al., 2013). So far, the only characterised SLiM for PP2A is the LxxIxE motif that binds to the regulatory subunit of the PP2A-B56 holoenzyme complex (Hertz et al., 2016).

This simplistic model of co-localisation driving function explains nicely how these phosphatases can target specific substrates, but it does not explain why these substrates select to interact specifically with one phosphatase over the other when their catalytic activities are apparently very similar. In that sense, it fails to capture the essence of why PP1 and PP2A have evolved to regulate different signals. They must presumably possess specific features that are repeatedly selected for by different pathways throughout the course of evolution, although exactly what these features are still remains unclear. It is important to address this because it may help to reveal why these two major phosphatase families have evolved to fulfil different signalling roles.

One major distinction between PP1 and PP2A is their ability to be regulated differently. This can occur directly on the holoenzymes, for example, via catalytic subunit phosphorylation or the binding of catalytic inhibitors (Grallert et al., 2015; Rogers et al., 2016; Verbinnen et al., 2017). A well-studied example of this is the inhibition of PP2A-B55 during mitosis by the ARPP19/ENSA phospho-proteins (Gharbi-Ayachi et al., 2010; Mochida et al., 2010). Whilst direct regulation of the holoenzyme is useful for modulating global phosphatase activity, there are many situations when individual pathways or substrates must be regulated separately. In these cases, the regulation can occur directly on the SLiMs within these pathways that are needed to direct the phosphatases towards specific substrates. Interestingly, in this respect, PP1 and PP2A-B56 behave in opposite ways: PP1 binding to the RVxF motif can be repressed by phosphorylation (Kim et al., 2003; Nasa et al., 2018), whereas PP2A-B56 interaction with the LxxIxE motif can be enhanced by phosphorylation (Hertz et al., 2016; Wang et al., 2016a; Wang et al., 2016b). These unique modes of phospho-regulation could allow PP1 and PP2A-B56 to perform very different signalling roles, however, it is difficult to dissociate whether it is these properties or others, such as catalytic preferences, that are more important in any given situation.

To investigate this further, we chose to focus on the kinetochore, which is a multi-complex structure assembled on chromosomes during mitosis to mediate their attachment to microtubules. Although this complex contains over 100 different proteins, PP1 and PP2A-B56 are recruited via their SLiMs to the same molecular scaffold - Knl1 - to regulate kinetochore-microtubule attachments and the spindle assembly checkpoint (SAC) (Saurin, 2018). These processes are critical for genome stability because microtubules bind to kinetochores to segregate the duplicated chromosomes equally and the SAC holds the mitotic state to give time for these microtubules to attach correctly. Importantly, even though PP1 and PP2A are recruited to a very similar molecular space on kinetochores, they still appear to control these key mitotic processes differently; as evidenced by the fact that removing either phosphatase produces markedly distinct phenotypic effects (these will be discussed in detail below) (Saurin, 2018). It is currently unclear how these phosphatases achieve specificity in such a crowded molecular environment, or indeed, why they are both needed to carry out different roles at the kinetochore. We therefore rationalised that this would be an ideal system to reveal answers about phosphatase specificity and functional diversity within the broader signalling context.

Using the direct approach of switching the phosphatases or their SLiMs at the kinetochore, we demonstrate that their unique phenotypic effects cannot be explained by either catalytic preferences or positional differences. Instead, we demonstrate that phenotypic diversity arises because the phosphatases are recruited via SLiMs that display opposite phospho-dependencies and, as a result, are subject to different forms of cross-regulation and feedback. Therefore, this study explains how downstream “specificities” can depend entirely on the mode of upstream regulation and it establishes a paradigm to explain how these two major phosphatase families may have evolved to couple to phosphorylation inputs in opposite ways.

## Results

### PP1-Knl1 and PP2A-B56 exert control over different kinetochore processes

Knl1 is a key signalling scaffold that functions at kinetochores to generate the SAC signal and regulate the attachment of microtubules. Critical for both of these processes are the ‘MELT’ motifs (for the consensus sequence: Met-Glu-Leu-Thr) that are scattered along the N-terminal half of Knl1 and phosphorylated by Mps1 kinase to recruit the Bub1/BubR1/Bub3 complex (London et al., 2012; Overlack et al., 2015; Primorac et al., 2013; Shepperd et al., 2012; Vleugel et al., 2013; Yamagishi et al., 2012; Zhang et al., 2014). This complex has two main functions: 1) it modulates Aurora B activity to regulate kinetochore-microtubule attachments (Aurora B is a kinase that can phosphorylate kinetochores to detach microtubule fibres (Krenn and Musacchio, 2015)), and 2) it provides a platform to recruit all other proteins needed for the SAC to delay mitotic exit (Saurin, 2018). Crucially, both of these functions are regulated by two co-localised phosphatase complexes: PP1 that is bound to SILK and RVSF motifs in the N-terminus of Knl1 (PP1-Knl1; note that Aurora B phosphorylates these motifs to inhibit PP1 binding) (Espeut et al., 2012; Liu et al., 2010; Meadows et al., 2011; Nijenhuis et al., 2014; Rosenberg et al., 2011) and PP2A-B56 that binds to an LSPIIE motif in BubR1 (note that Cdk1 and Plk1 both phosphorylate this motif to enhance PP2A-B56 interaction) (Espert et al., 2014; Foley et al., 2011; Kruse et al., 2013; Nijenhuis et al., 2014; Suijkerbuijk et al., 2012; Xu et al., 2013). There has been debate surrounding exactly which phosphatase controls what process (Saurin, 2018), therefore we begin by carefully dissecting their individual roles at the kinetochore.

As shown previously by others (Shrestha et al., 2017), removal of the PP2A-B56 SLiM in BubR1 (BubR1^ΔPP2A^) causes severe defects in chromosome alignment, whereas inactivation of the PP1 SLiM in Knl1 (Knl1^ΔPP1^) does not (fig.1a). These defects are associated with enhanced phosphorylation of the Ndc80 tail region (fig.1b and supp.fig.1a); a key Aurora B substrate that must be dephosphorylated to stabilise kinetochore-microtubule attachments (Krenn and Musacchio, 2015). In contrast to these differential effects on chromosome alignment, PP1-Knl1 or PP2A-B56 are both needed to allow Knl1 MELT dephosphorylation and SAC silencing following inhibition of the upstream kinase Mps1 (fig.1c,d and supp.fig.1b,c) (Espert et al., 2014; Nijenhuis et al., 2014). However, even in this situation, the BubR1^ΔPP2A^ and Knl1^ΔPP1^ phenotypes differ because the effects of PP2A-B56 loss can be specifically rescued by Aurora B inhibition (fig.1e,f and supp.fig.1d,e; note that this is not due to differential effects on microtubule attachments because all SAC assays were performed in nocodazole to depolymerise microtubules). We hypothesised previously that PP2A-B56 sits upstream of PP1 in SAC silencing by supressing Aurora B-mediated phosphorylation of Knl1 to allow PP1-Knl1 association (Nijenhuis et al., 2014). This is consistent with the results in figure 1g, which show that mutating these Aurora B sites in Knl1 (Knl1^PP1(2A)^) allows nocodazole-arrested BubR1^ΔPP2A^ cells to exit mitosis rapidly following Mps1 inhibition. Therefore, rescuing PP1-Knl1 can bypass the requirement for PP2A-B56 in SAC silencing. Importantly, the same is not true in reverse, because PP2A-B56 is present on kinetochores in Knl1^ΔPP1^ cells (fig.1h and supp.fig.1f), and yet these cells can still not silence the SAC (fig.1d) (Nijenhuis et al., 2014).

**Figure 1.**
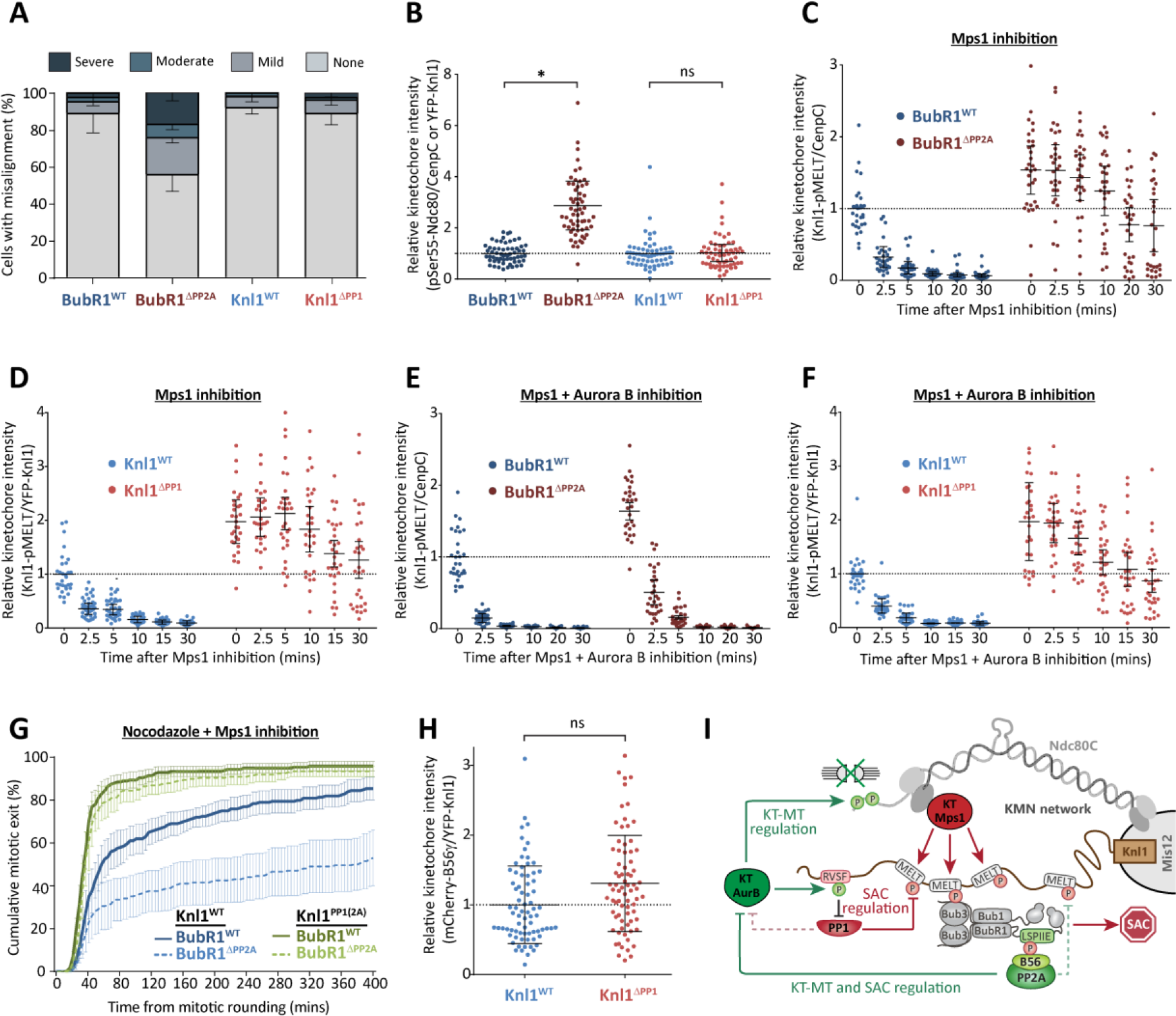
PP1-Knl1 and PP2A-B56 exert control over different pathways and processes at the kinetochore. **A,B.** Effect of phosphatase binding mutants on chromosomal alignment (**A**) and pSer55-Ndc80 kinetochore levels in nocodazole (**B**). Graphs in A shows the mean (-SD) of 3 experiments, 100 cells per condition per experiment. Graphs in B show data from 60 cells from 4 experiments. **C-F.** Effect of phosphatase binding mutants on Knl1-MELT dephosphorylation in nocodazole-arrested cells treated with the Mps1 inhibitor AZ-3146 (2.5μM) for the indicated times; either alone (**C,E**) or in combination with the Aurora B inhibitor ZM-447439 (2μM; **D,F**). MG132 was included in all treatments to prevent Cyclin B degradation and mitotic exit following Mps1 inhibition. **G**. Duration of mitotic arrest in cells, expressing various wild type and phosphatase-binding mutant combinations, treated with nocodazole and 2.5 µM AZ-3146. Graph shows cumulative mean (±SEM) of 3 experiments, 50 cells per condition per experiment. **H.** Kinetochore B56γ levels in nocodazole-arrested cells expressing wild type or PP1-binding deficient Knl1 (Knl1^ΔPP1^). Graph shows 75 cells per condition from 4 experiments. **I.** A schematic model to depict the primary effects of PP1 and PP2A-B56 at the KMN network. Note that PP2A-B56 is shown to regulate Aurora B directly, but this is simply meant to represent co-antagonism of both Aurora B substrates at the kinetochore (pRVSF and pNdc80). For all kinetochore intensity graphs, each dot represents a cell and the error bars display the variation between the experimental repeats (displayed as ±SD of the experimental means). ns p>0.05, *p<0.05.

In summary, PP1 and PP2A-B56 are recruited via their respective SLiMs to a very similar molecular space on kinetochores and yet they still manage to exert control over different substrates and different mitotic processes. PP2A-B56 antagonises Aurora B signals to regulate kinetochore-microtubule attachments and PP1-Knl1 interaction, whereas PP1-Knl1 antagonises Mps1 signals to silence the SAC (fig.1i). PP2A-B56 may also contribute to SAC silencing directly, but it cannot support MELT dephosphorylation without PP1-Knl1 (fig.1d,h). Similarly, PP1-Knl1 could help to stabilise initial microtubule attachments, but it is not essential, and it is also not able to support proper chromosome alignment in the absence of PP2A-B56 (fig.1a). Therefore, both of these potential links are still included in the model in Figure 1i, but only as dotted arrows.

### PP1 and PP2A-B56 can functionally substitute for each other at kinetochores

The simplest explanation for the observed phenotypic differences is that PP1 and PP2A are different catalytic enzymes that could produce specific effects at the kinetochore. Therefore, to test whether the identity of the phosphatase is a key determinant of their function, we deleted the SLiMs that recruit PP1 or PP2A-B56 to kinetochores and asked whether the resulting phenotypes could be rescued if the alternative phosphatase is recruited in its place. Figure 2a shows that the chromosome alignment defects following truncation of BubR1 before the PP2A-B56 binding region (BubR1^ΔCT^), can be rescued if a short region from the Knl1 N-terminus is fused in its place to recruit PP1 (BubR1^ΔCT-PP1:Knl1^; note that the Aurora B sites are mutated in the SLiMs to prevent Aurora B from inhibiting PP1 binding). This is dependent on direct PP1 recruitment because the effect is lost if the N-terminal fusion has the PP1-binding SLiM mutated (BubR1^ΔCT-Knl1^). Conversely, if the first 70 amino acids of the Knl1 N-terminus are removed, which contains the PP1 binding region (Knl1^ΔNT^), then SAC silencing and MELT dephosphorylation are inhibited following Mps1 inhibition (fig.2b,c and supp.fig.2a). However, if B56 is tethered directly to the N-terminus of Knl1 (B56γ-Knl1^ΔNT^) then both of these effects can be fully rescued (fig.2b,c and supp.fig.2a). This requires PP2A catalytic activity, because fusion of a B56 mutant that cannot bind the catalytic domain (B56γ^CD^-Knl1^ΔNT^ (Vallardi et al., 2018)) does not support SAC silencing. Finally, preventing PP2A-B56 recruitment to BubR1 also gives a SAC silencing defect, and this can also be fully rescued by recruiting PP1 in its place (fig.2d,e and supp.fig.2b). Therefore, both phosphatases can functionally substitute for each other if their respective positions at the kinetochore are switched. This demonstrates that the phenotypic differences cannot be explained by the identity of the individual phosphatases.

**Figure 2.**
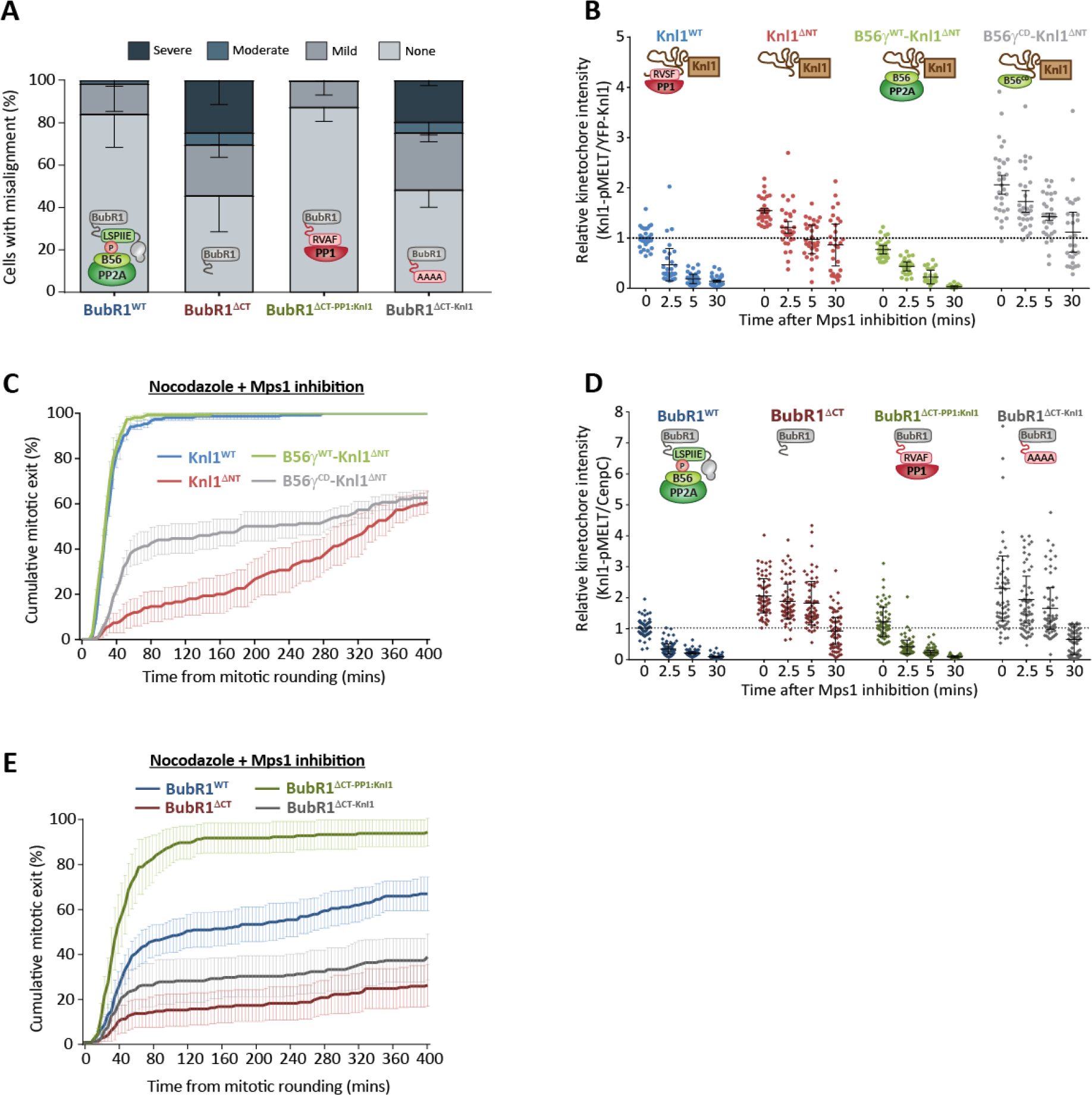
PP1 and PP2A-B56 can functionally substitute for each other at the kinetochore. **A.** Effect of altering the phosphatase at BubRl on chromosomal alignment. Graph shows mean (-SD) of 3 experiments, with at least 100 cells per condition per experiment. **B,C.** Effect of altering the phosphatase at Knl1 on SAC phenotypes. **B.** Knl1-MELT dephosphorylation in nocodazole-arrested cells treated with the Mps1 inhibitor AZ-3146 (2.5μΜ) for the indicated times. Graph shows 30 cells per condition from 3 experiments. **C.** Duration of mitotic arrest in cells treated with nocodazole and 5 μΜ AZ-3146. Graphs shows cumulative mean (±SEM) of 3 experiments, 50 cells per condition per experiment. **D,E**. Effect of altering the phosphatase at BubR1 on SAC phenotypes. **D.** Knl1-MELT dephosphorylation in nocodazole-arrested cells treated with the Mps1 inhibitor AZ-3146 (2.5μΜ) for the indicated times. Graph shows 50 cells per condition from 6 experiments. **E.** Duration of mitotic arrest in cells treated with nocodazole and 2.5 μΜ AZ-3146. Graph shows cumulative mean (±SEM) of 5 experiments, 50 cells per condition per experiment. MG132 was included in treatments in B and D to prevent Cyclin B degradation and mitotic exit following Mps1 inhibition. For all kinetochore intensity graphs, each dot represents a cell and the errors bars display the variation between the experimental repeats (displayed as ±SD of the experimental means).

### PP1 and PP2A-B56 can function from different positions at the KMN network

If identity is not important for function, then the precise positions may be critical instead. For example, although PP1 or PP2A are recruited to the same molecular subcomplex on kinetochores, they may only have restricted access to a subset of different substrates from their exact positions on Knl1 and BubR1. To address this, we first focussed on kinetochore-microtubule attachment regulation because this was clearly defective when phosphatases were absent from the BubR1 position (figs 1a and 2a). Importantly, however, this position does not appear to be critical, because chromosomal alignment defects in BubR1^ΔPP2A^ cells could be rescued by recruiting B56 to the N-terminus of Knl1 instead (B56γ-Knl1^ΔNT^) (fig.3a). Therefore, PP2A-B56 can support chromosomal alignment from either the BubR1 or Knl1 position.

**Figure 3.**
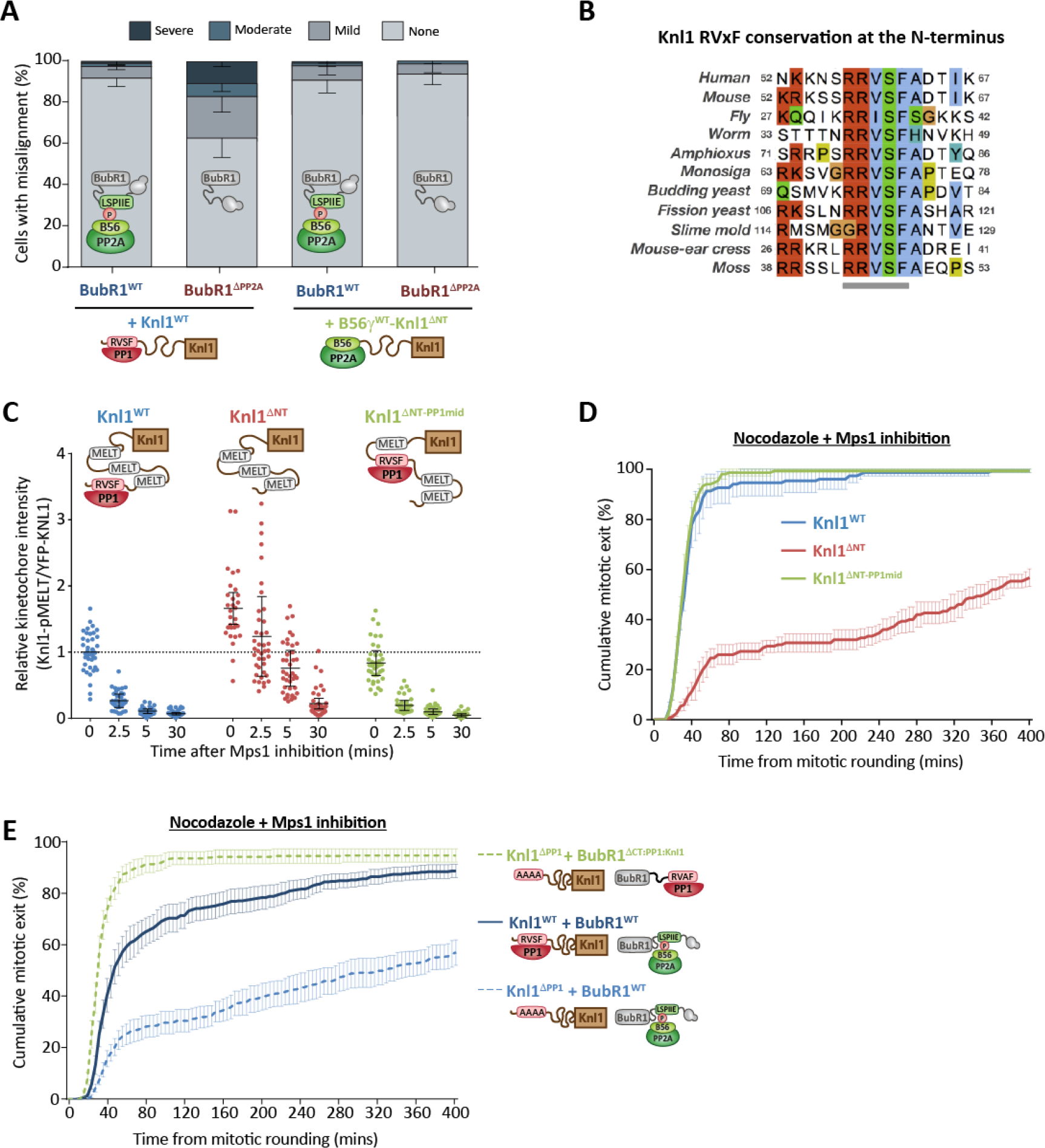
The exact positions of PP1 and PP2A-B56 are not critical for their kinetochore functions. **A**. Chromosomal alignment after removing PP2A-B56 from BubR1 and repositioning it at Knl1. Graph shows the mean (-SD) of 3 experiments, 100 cells per condition per experiment. **B.** Conservation of the RVxF SLIM at the N-terminus of Knl1. Sequences are coloured using ClustalW scheme. **C,D**. Effect of relocating the PP1 SLiMs to the middle of Knl1 on SAC phenotypes. **C.** Knl1-MELT dephosphorylation in nocodazole-arrested cells treated with the Mps1 inhibitor AZ-3146 (5μM) for the indicated times. Graph shows 40 cells per condition from 4 experiments. MG132 was included to prevent Cyclin B degradation and mitotic exit following Mps1 inhibition. Each dot represents a cell and the errors bars display the variation between the experimental repeats (displayed as ±SD of the experimental means). **D.** Duration of mitotic arrest in cells treated with nocodazole and 5 μΜ AZ-3146. **E.** Effect of switching PP1 from Knl1 to BubR1 on SAC silencing. Duration of mitotic arrest in cells, expressing different and wild type mutant combinations, treated with nocodazole and 2.5 μΜ AZ-3146. Graph in D and E show the cumulative mean (±SEM) of 3 experiments, 50 cells per condition per experiment. ns p>0.05, *p<0.05.

We next examined why PP1-Knl1 was sufficient on its own to support SAC silencing, whereas PP2A-B56 bound to BubR1 was not (fig.1c-h). The SLiM that recruits PP1 is conserved at the N-terminus of Knl1 throughout evolution (fig.3b and supp.fig.3a), therefore we hypothesized that this position may be critical to access the MELTs. Surprisingly, however, moving the PP1-binding SLiM into the middle of Knl1 (Knl1^ΔNT-PP1mid^), had little effect on MELT dephosphorylation (fig.3c, supp.fig.3b), Bub complex removal from kinetochores (supp.fig.3c-f) or SAC silencing (fig.3d) following Mps1 inhibition in nocodazole. Therefore, PP1 does not need to be positioned at the N-terminus of Knl1 to silence the SAC in the absence of microtubules. Although the exact position does not appear to be important, the phosphatase could still require a unique feature of Knl1 itself, such as its predicted flexibility, to silence the SAC. This might explain why PP2A-B56 bound to BubR1 could not dephosphorylate the MELT motifs in Knl1^ΔPP1^ cells (fig.1d,h). Although we had already observed that PP1 can silence the SAC when bound to BubR1 (fig.2d,e), this effect might be direct or indirect, because PP1 could simply dephosphorylate the SILK/RVSF motifs to recruit an additional PP1 molecule to Knl1. To distinguish between these possibilities, we created double mutant cells in which the BubR1 phosphatase could be switched in either the presence or absence of PP1-Knl1. Importantly, SAC silencing was still rescued in Knl1^ΔPP1^ cells by a BubR1 mutant that recruits PP1, even though it could not be recovered in the same cells by a BubR1 wild type that can bind to PP2A-B56 (fig.3e). Therefore, PP1 can silence the SAC directly when positioned at either Knl1 or BubR1.

In summary, PP1 and PP2A-B56 have specific functions at the kinetochore (fig.1), but these cannot be explained by differences in either their catalytic subunits (fig.2) or their spatial positioning (fig.3). This is surprising because these are thought to be the principal determinants of phosphatase specificity, and if these phosphatases don’t display any obvious ‘specificity’ then it is not easy to rationalise why they produce differential effects at the kinetochore. Furthermore, if the identity of the phosphatase is not important, as fig.2 demonstrates, then it is not clear why there is such a difference in the ability of PP1 or PP2A to support kinetochore-microtubule attachment from the Knl1 N-terminus (compare both BubR1^ΔPP2A^ conditions in fig.3a) and SAC silencing from the BubR1 position (compare both Knl1^ΔPP1^ conditions in fig.3e). However, as well as switching phosphatases in these key experiments, we also abolished their regulation by phosphorylation inputs. In particular, Aurora B phosphorylates the Knl1-SLiMs to inhibit PP1 (Liu et al., 2010), but when microtubule attachments were rescued by the recruitment of PP2A-B56 to Knl1, we directly tethered B56 and lost these regulatory inputs (fig.3a). In addition, Cdk1 and Plk1 phosphorylate the BubR1-SLiM to recruit PP2A-B56 (Elowe et al., 2007; Huang et al., 2008; Kruse et al., 2013; Suijkerbuijk et al., 2012; Wang et al., 2016a; Wang et al., 2016b), but when the SAC was rescued by recruiting PP1 to BubR1 we removed this phospho-dependence (fig.3e). Therefore, we rationalised that it may be the unique forms of SLiM regulation that prevents PP1-Knl1 from stabilising microtubule attachments when PP2A-B56 is removed, and restricts PP2A-B56 from silencing the SAC when PP1-Knl1 is absent.

### PP1-Knl1 and PP2A-B56 use opposite phospho-dependencies to control distinct kinetochore processes

A major difference in their SLiM regulation is that phosphorylation of Knl1 represses PP1 binding, whereas phosphorylation of BubR1 enhances PP2A-B56 binding. Therefore, even if these phosphatases display no downstream specificity at kinetochores, removal of their SLiMs will enhance phosphorylation of the opposing SLiM and produce opposite effects on phosphatase localisation (fig.4a). Indeed, inhibiting BubR1:PP2A-B56 interaction is known to enhance Aurora B-mediated phosphorylation of the Knl1 SLiM to prevent PP1 binding (Nijenhuis et al., 2014). This is not a specific effect of PP2A-B56 *per se*, because removal of PP2A-B56 from BubR1 enhances Knl1-RVSF phosphorylation, and this can be rescued by recruiting PP1 to BubR1 instead (fig.4b and supp.fig.4a). Therefore, inhibiting phosphatase activity at BubR1 also inhibits it at Knl1, because the PP1:SLiM interaction is repressed by phosphorylation. Importantly, if these phosphorylation sites are mutated to alanine to rescue PP1-Knl1 in BubR1^ΔPP2A^ cells (Knl1^PP1(2A)^), then chromosome alignment defects are also recovered (fig.4c). Therefore, either phosphatase in either position can support chromosomal alignment (figs.2a, 3a and 4c). There appears to be a specific role for PP2A-B56 because when it is removed from BubR1 then PP1-Knl1 is also lost (assuming Aurora B is active on kinetochores to phosphorylate the Knl1 SLiMs). In contrast, PP1-Knl1 is redundant because even when this is removed, PP2A-B56 is still preserved on kinetochores to antagonise Aurora B (fig.1h). Therefore, the different forms of SLiM interaction explain why chromosomal alignment is primarily controlled by distinct phosphatase complexes at kinetochores (fig.1a).

**Figure 4.**
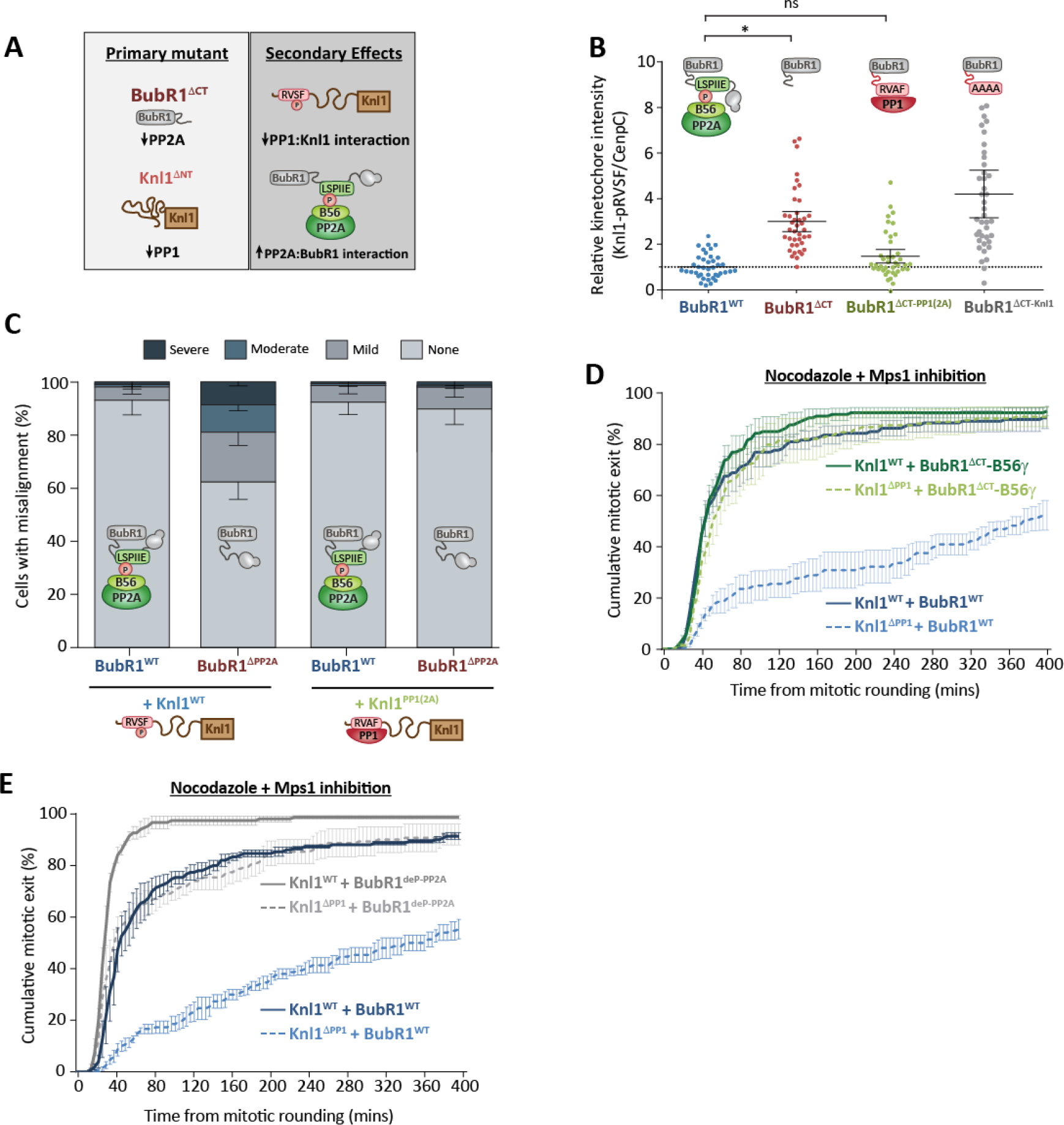
PP1-Knl1 and PP2A-B56 use opposite phospho-dependencies to control distinct processes at the kinetochore. **A.** Schematic to illustrate how cross-regulation between SLiMs affects kinetochore phosphatase levels. **B.** Effect of altering the phosphatase at BubR1 on Knl1-pRVSF kinetochore levels in nocodazole. Graph shows data from 30 cells from 3 experiments. Each dot represents a cell and the error bars display the variation between the experimental repeats (displayed as ±SD of the experimental means). **C.** Chromosome alignment in cells expressing mutant combinations to prevent phosphorylation of the PP1 SLiMs following removal of PP2A-B56 from BubR1. Graphs shows the mean (-SD) of 3 experiments, 100 cells per condition per experiment. **D,E.** Effect of removing the phospho-dependence of PP2A-B56 on SAC silencing. Duration of mitotic arrest in cells, expressing various wild type and mutant combinations, treated with nocodazole and 2.5 μM AZ-3146. Graphs show the cumulative mean (±SEM) of 3 experiments, 50 cells per condition per experiment.

To determine the reason for the differential effect on the SAC, we focussed on the crucial observation that PP1-Knl1 inhibition prevents SAC silencing even though PP2A-B56 remains bound to BubR1 at kinetochores (fig.1d,h). We hypothesised that the phospho-dependence of this BubR1 interaction restricts PP2A-B56 from efficiently silencing the SAC, which is supported by the observation that PP1 can silence the SAC when recruited to BubR1 in a manner that is independent of phosphorylation (BubR1^ΔCT-PP1:Knl1^ in fig.3e). In agreement with this hypothesis, a similar effect is also observed if B56 is fused directly to BubR1 (BubR1^ΔCT^-B56γ in fig.4d), which demonstrates that PP1 and PP2A-B56 can both silence the SAC efficiently when tethered directly to BubR1; even in Knl1^ΔPP1^ cells. These fusions eliminate the dependence on phosphorylation for phosphatase recruitment, but in addition, they also change the relative orientation of the phosphatases at BubR1. This could, in principle, provide the additional flexibility needed for access to key substrates that might otherwise be inaccessible when B56 is bound to the LxxIxE motif. Increased flexibility is unlikely to explain why PP2A-B56 cannot silence the SAC, however, given that insertion of a flexible linker immediately before the PP2A binding motif in BubR1 does not affect MELT dephosphorylation or SAC silencing in either the presence or absence of PP1-Knl1 (supp.fig.5). Nevertheless, to test directly whether a lack of phospho-dependence was the reason for enhanced SAC silencing, we mutated the PP2A binding sequence in BubR1 to an LxxIxE sequence that binds to B56 in the same manner and with similar affinities, but, crucially, does not depend on phosphorylation (BubR1^deP-PP2A^ which uses an LPTIHE sequence (Kruse et al., 2018)). Fig.4e shows that BubR1^deP-PP2A^ cells were now able to silence the SAC in the absence of PP1-Knl1, demonstrating that PP2A-B56 is restricted from silencing the SAC due to a phospho-dependent interaction with BubR1. There is still an additional contribution of PP1-Knl1 in this situation (compare BubR1^deP-PP2A^ in Knl1^WT^ and Knl1^ΔPP1^ cells), which likely indicates that both phosphatases collaborate to shut down the SAC. This is predicted given that both phosphatases are indistinguishable in our assays if they are coupled in either position independently of phosphorylation (figs.2b-e).

In summary, although PP1 and PP2A-B56 are indistinguishable in our assays when their positions are switched (fig.2), they can still produce distinct effects because they couple to phosphorylation inputs in opposite ways (fig.4). We therefore next sought to address whether this alone was sufficient to explain their phenotypic differences at kinetochores. To this end, we developed a mathematical model of the network outlined in fig.1i. A crucial aspect of this model, which is displayed schematically in fig.5a, is that both phosphatases dephosphorylate the same substrates (Knl1-pMELT, Knl1-pRVSF, BubR1-pLSPI and pNdc80) with identical kinetics when docked to their native SLiMs on Knl1. This binding occurs directly for PP1 (via dephospho-Knl1-RVSF) or indirectly for PP2A (via phospho-Knl1-MELT and phospho-BubR1-LSPI). The kinases that phosphorylate these docking motifs (Mps1, Aurora B, Cdk1) are given a fixed activity that is not regulated by the phosphatases or any other aspect of the model. Therefore, any difference between the two phosphatases in the model is due to their inverse phospho-dependencies, as suggested by all results presented thus far.

**Figure 5.**
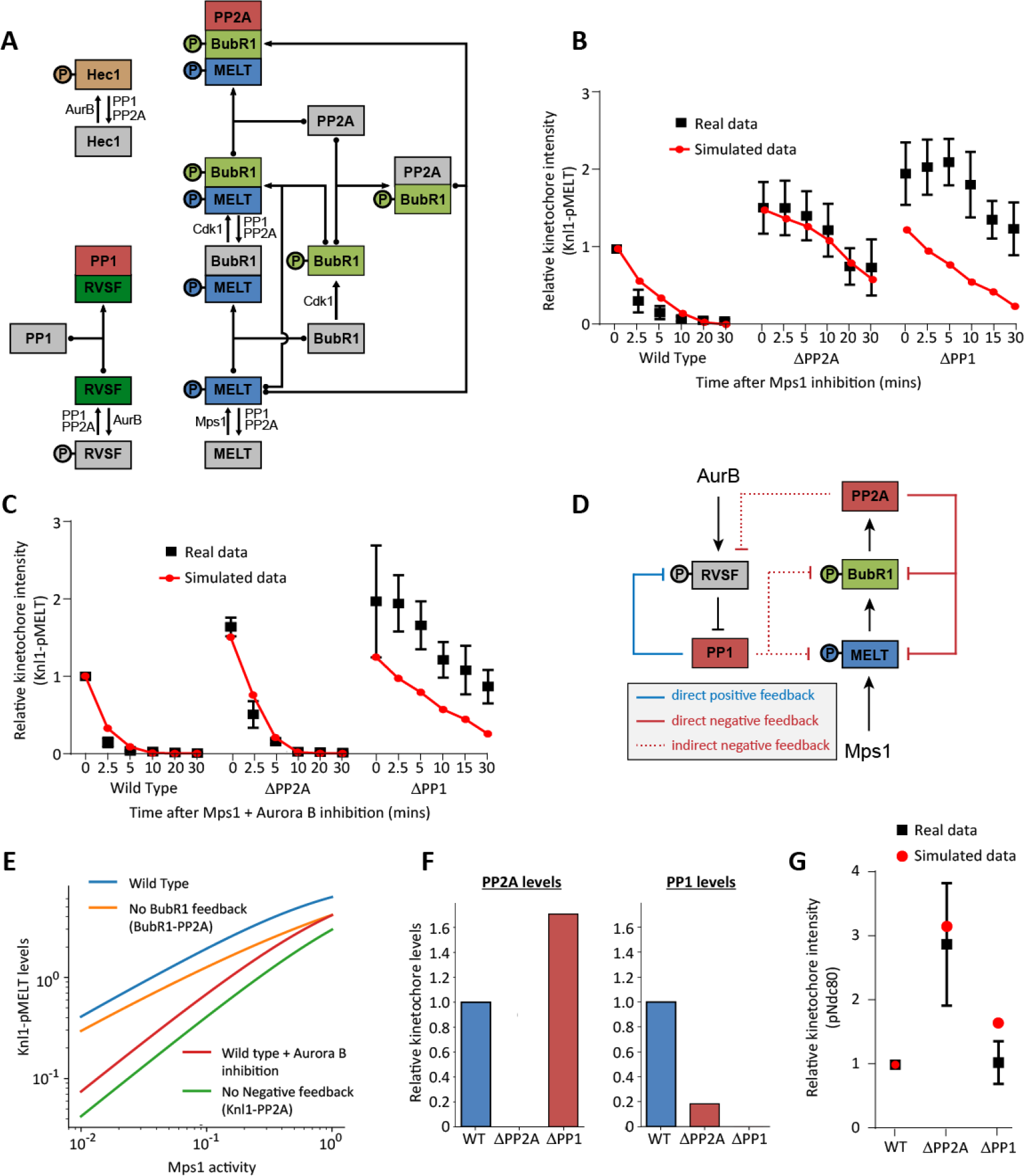
Mathematical model to show how identical phosphatases with opposite phospho-dependencies can produce distinct phenotypic behaviours. **A.** Full wiring diagram underlying the mathematical model that assumes identical activities of PP1 and PP2A towards all kinetochore substrates. Their only differences are their opposite modes of phospho-regulation. Arrows with dotted ends represent reversible binding/dissociation reactions. Regular arrows represent phosphorylation/dephosphorylation reactions catalysed by the kinases/phosphatases, as indicated. **B,C.** Comparison of the simulated output with the real data from figure 1c-d. Graphs show Knl1-MELT dephosphorylation after Mps1 inhibition (**B**) or Mps1 + Aurora B inhibition (**C**). **D.** Schematic to illustrate the various positive and negative feedback loops that affect phosphatase recruitment. Note that both phosphatases act on all phosphorylation sites **E.** Simulated steady state levels of Knl1-pMELT as a function of Mps1 for different conditions to remove negative feedback or inhibit Aurora B. Sensitivity/robustness can be directly compared because it corresponds to the slope of the curve on the log-scale graph. **F.** Simulated kinetochore levels of PP1 or PP2A after removal of either phosphatase. The simulated output confirms predictions for cross-regulation in figure 4a. **G.** Comparison of simulated pNdc80 steady state values during a SAC arrest (i.e. Mps1 and Aurora B active) to the experimental data from figure 1b.

### Modelling to show how identical phosphatases with opposite phospho-dependencies can produce distinct phenotypic behaviours

We first asked whether the model could reproduce the SAC data from figures 1c-f. In our simulations, Knl1-MELT is dephosphorylated rapidly upon Mps1 inhibition and this is dependent on the presence of both PP1 and PP2A. Combined Aurora B inhibition, speeds up the rate of MELT dephosphorylation and specifically rescues the effects of PP2A loss (fig.5b,c). Therefore, the model can reproduce the core data, which crucially, also includes one key unexplained aspect of our results: PP2A-B56 is unable to silence the SAC in the absence of PP1. We were able to explore this further in the model to demonstrate that negative feedback loops downstream of PP2A prevent this phosphatase from silencing the SAC efficiently. Negative feedback occurs at multiple levels because PP2A dephosphorylates both BubR1 and Knl1 to effectively remove and inhibit its own recruiting SLiM (this occurs directly and indirectly via PP1; see fig.5d). We had already determined how much of an effect these feedback loops have on PP2A-B56 localisation, by quantifying kinetochore B56γ levels following expression of a wild type or inactive B56 mutant (B56γ^CD^) (supp.fig.6) and then including this difference in the model. Now, by selectively removing the loops we can measure their effects on the output. This demonstrates that preventing the feedback onto phospho-BubR1 reduces phospho-MELT levels (fig.5e), which is consistent with the fact that PP2A can silence the SAC effectively in BubR1-B56 or BubR1^deP-PP2A^ cells (figs.4d,e). However, a stronger effect on MELT dephosphorylation is observed when all negative feedback loops are abolished by recruiting a phosphatase that is independent of either phospho-BubR1 or phospho-Knl1 (fig.5e). In this case there is also a significantly increased sensitivity to changes in Mps1 levels. This is effectively the situation that is achieved when Aurora B is inhibited and PP1-Knl1 recruitment to kinetochores is unconstrained, as can be seen in the WT situation with Aurora B inhibition (fig.5e).

Therefore, the model illustrates that negative feedback downstream of PP2A can allow the SAC to remain robust to variations in Mps1 activity (i.e. Knl1-pMELT levels remain high when Mps1 levels decrease; Fig.5e) by limiting the ability of this phosphatase to dephosphorylate the MELTs on its own. When Aurora B activity falls at kinetochores, then PP1 recruitment is elevated and the SAC can be silenced efficiently without the effects of negative feedback restricting phosphatase levels. In fact, this transition is aided by positive feedback instead, because PP1 dephosphorylates the KNl1-RVSF motif to enhance its own recruitment (fig.5d). Using identical parameters to the SAC simulation, the model also simulates the cross-regulation that is illustrated in figure 4a. In the absence of PP2A, PP1 levels are dramatically reduced, whereas in the absence of PP1, PP2A levels of increased (fig.5f). This is consistent with the data from figure 1h and leads to a differential effect on steady state pNdc80 levels (fig.5g). This is also consistent with our observed differential effects on kinetochore-microtubule attachments and Ndc80 phosphorylation (fig.1a,b).

The simulation therefore illustrates how identical phosphatases can produce differential phenotypic behaviours by using opposite modes of phospho-regulation. Considering we observe no other differences between PP1 and PP2A-B56 in any of our assays, this implies that kinetochores have evolved to interact with these phosphatases primarily because of their inverse phospho-dependencies. This has important implications for signalling in general, because it is likely that many other pathways also select for these key defining features.

### Phospho-regulation is a common feature of RVxF and LxxIxE SLiMs

To analyse this further, we curated a list of validated and predicted RVxF and LxxIxE motifs, which are present in almost 700 unique proteins (supp.table.1). Motif analysis demonstrates that serines and threonines are statistically enriched at positions within each motif where phosphorylation is known to inhibit (RVxF) or enhance (LxxIxE) phosphatase interaction (fig.6a,b) (Hertz et al., 2016; Kim et al., 2003; Kumar et al., 2016; Nasa et al., 2018; Wang et al., 2016a; Wang et al., 2016b). Furthermore, up to 25% of the validated motifs are known to be phosphorylated in vivo, and 50% of the RVxF and 100% of the LxxIxE motifs contain phosphorylatable residues at the key positions (fig.6c-e); which is a statistically significant enrichment (see amino acid matrices in supp.table.1). It should be noted that phosphorylation of residues outside of the core RVxF region can also inhibit PP1 binding (Kumar et al., 2016; Qian et al., 2015). Furthermore, the negatively charged surface that surrounds the RVxF pocket on PP1 (fig.6f), could potentially mediate many other electrostatic interactions that are inhibitable by phosphorylation. Therefore, although only half of the core RVxF motifs contain phosphorylatable residues, the percentage that are phospho-regulatable is probably much higher. In contrast to PP1, the interaction between PP2A-B56 and LxxIxE motifs can be enhanced by phosphorylation inside and immediately after the core motif. This is because the charged phosphate residues in the P2 and P7-9 positions can make key electrostatic interactions with basic residues in a groove on B56 (Hertz et al., 2016; Wang et al., 2016a; Wang et al., 2016b) (fig.6g). Therefore, the binding pockets on PP1 and B56 appear to have evolved to respond to phosphorylated SLiMs in opposite ways and numerous pathways have taken advantage of these unique properties to enable localised phosphatase activity to be modulated by kinase inputs. This study therefore defines how two of the main phosphatase families in eukaryotic cells have evolved to perform very important, but also very distinct, signalling roles.

**Figure 6.**
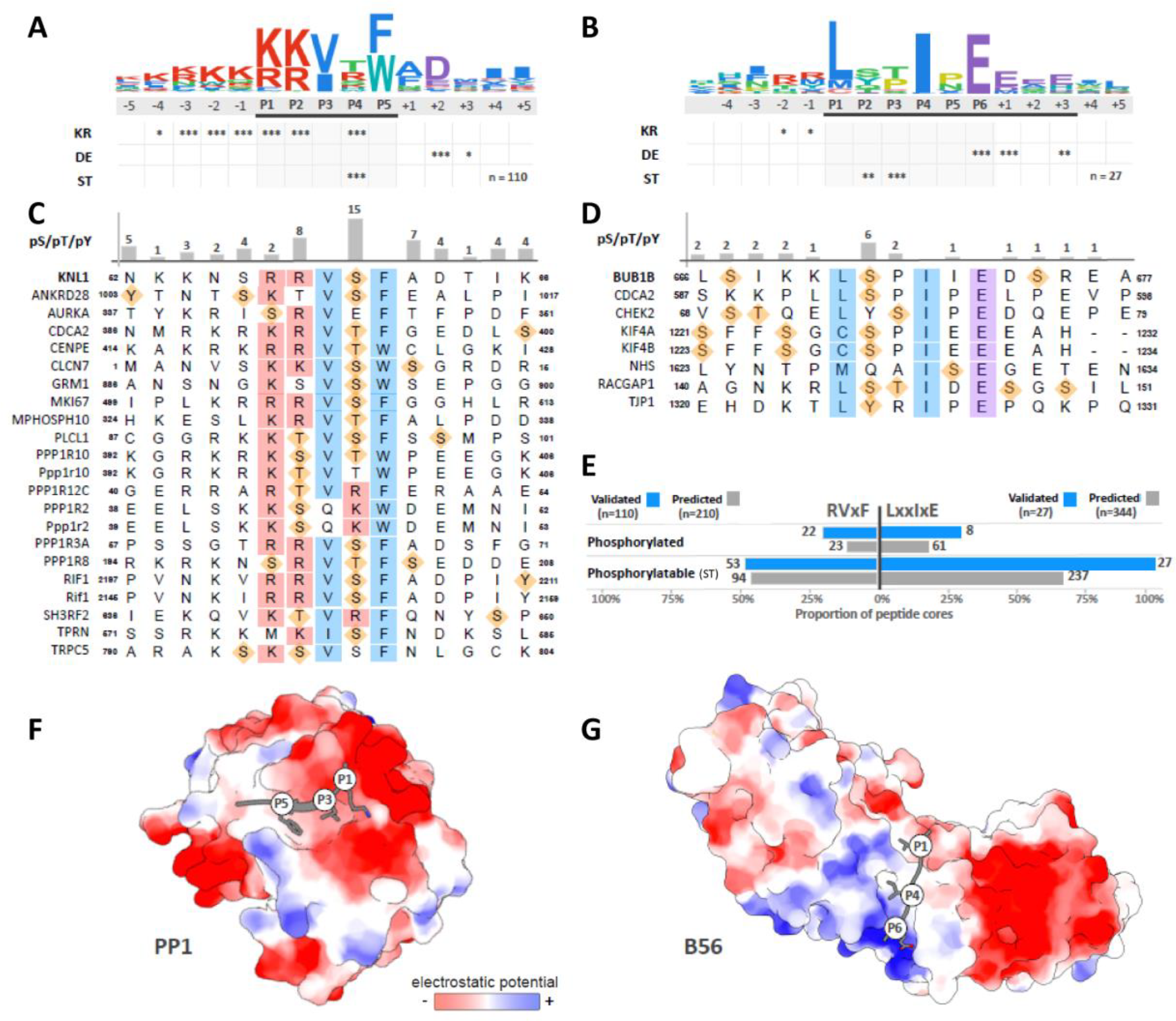
Analysis of the specificity determinants and phosphorylation sites in PP1 RVxF and PP2A-B56 LxxIxE motifs. **A.** A log^10^ relative binomial sequence logo based on 110 validated PP1-binding RVxF motifs. Asterisks denote the significance of the amino acid enrichment of basic (KR), acidic (DE) and phosphorylatable by Ser/Thr kinases (ST) (**\*** p **<** 0.01**, \*\*** p **<** 0.001**, \*\*\*** p **<** 0.0001). The logo is coloured using ClustalW colouring. **B.** As panel A, built using 27 validated PP2A-B56-binding LxxIxE motifs. **C.** Sequence of the 22 RVxF motifs with experimentally validated phosphorylation sites within the region of the motif (region defined by black line under the logo in panel A). Consensus positions are indicated by boxes coloured according to the ClustalW colouring scheme. Phosphorylated sites are indicated with orange diamonds. **D.** As panel C, but for the 8 LxxIxE motifs with experimentally validated phosphorylation sites. **E.** Summary of the LxxIxE and RVxF motifs phosphorylated (experimentally validated) and phosphorylatable (ST) within the motif regions (defined by black line under the logo in panels A/B). Data is shown for validated motifs (blue bars) and a set of high confidence predicted motifs (grey bars) created using the PSSMSearch software (Krystkowiak et al., 2018) by using the validated motifs as input and filtering as described in (Hertz et al., 2016). **F.** Structure of the RVxF motif of Rb bound to PP1 showing the key side chains of the motif that interact with the binding pocket (PDB ID:3N5U) (Hirschi et al., 2010). **G.** Structure of the LxxIxE motif of BubR1 bound to B56 showing the key side chains of the motif that interact with the binding pocket (PDB ID:5JJA) (Wang et al., 2016a). Structures are rendered using the Coulombic Surface Colouring in the Chimera package to show surface charge around the motif binding pockets.

## Discussion

The integration of kinase and phosphatase signals is a critical aspect of signal transduction (Gelens et al., 2018). At the kinetochore, numerous kinase and phosphatase signals converge at the KMN network to regulate two key mitotic process (Saurin, 2018). We show here that the two distinct phosphatases in this case are used for their ability to positively or negatively respond to kinase inputs. Although the kinases themselves clearly play an important role in determining phosphatase function, as will be discussed further below, it is important to point out that the differential effects illustrated here are primarily caused by cross-regulation and feedback between phosphatases.

The cross-regulation occurs because the phosphatases are embedded in the same network and can therefore dephosphorylate the respective SLiMs to produce opposing effects on each other’s recruitment (fig.4a and 5f). In particular, PP2A can dephosphorylate the SILK and RVSF SLiMs to enhance PP1-Knl1 levels and exert control over net phosphatase activity at the KMN network. This control is relinquished upon loss of Aurora B activity, because these SLiMs are no longer phosphorylated, which ensures that PP1-Knl1 can then be recruited independently of PP2A-B56. This is likely to be important to allow SAC silencing and microtubule stability to be maintained when PP2A-B56 is removed from kinetochores under tension.

In addition to cross-regulation, their unique phospho-dependencies also elicit different forms of feedback regulation. There are many feedback loops to consider (fig.5d), but the underlying theme is that phosphatase activity helps to enhance PP1 and repress PP2A. As a consequence, PP2A is subjected to a variety of negative feedback loops, which our modelling suggests, restricts this phosphatase from dephosphorylating the MELT motifs on its own following Mps1 inhibition. This contrasts with PP1, which engages in positive feedback and can dephosphorylate the MELT motifs in a more responsive manner (fig.5e). Aurora B activity safeguards the transition from PP2A to PP1 at kinetochores, which illustrates why it plays a key role in determining whether the SAC is robust or responsive to declining Mps1 activity. This has two important implications: 1) tension is ultimately required to inhibit Aurora B and allow the SAC to be silenced efficiently. This may explain why the SAC remains active on mono-orientated attachments and why it still takes hours to exit mitosis when stable attachments are formed that cannot generate tension (Etemad and Kops, 2016; Tauchman et al., 2015); 2) Aurora B is a potentially dangerous node in the network that could be hijacked by cancer cells to weaken the SAC and generate hyper-stable kinetochore-microtubule attachments (Cordeiro et al., 2018). These two effects can collaborate to generate high levels of chromosomal instability, therefore It will be important in future to determine whether Aurora B activity is commonly deregulated in cancer cells.

Another important implication of this work is that kinetochore-microtubule attachments and the SAC are principally regulated by different phosphatase complexes. We show that, despite their lack of specificity, PP2A-B56 is essential to stabilise microtubules attachments, but PP1-Knl1 is then ultimately required to help shut down the SAC. A similar separation of function was recently demonstrated in yeast, but in this case between different PP1 complexes (Roy et al., 2018). The general principle is the same in both cases, however, because sequential regulation by different phosphatases is predicted to guard against inappropriate SAC silencing when microtubule attachments are not correct.

The work presented here implies that kinetochores have evolved to interact with PP1 and PP2A-B56 mainly because of their opposite phospho-dependencies. It is important to clarify, however, that although these phosphatases were fully interchangeable in all our experiments, that does not exclude the possibility that important differences exist that were simply not detected in our assays. The relative activities of each phosphatase towards key phosphorylation sites on Knl1 look to be identical (fig.2b,d and fig,4b), but there are probably other important substrates that remain to be measured. Furthermore, there are other established regulatory mechanisms that do not involve the SLiMs, but which could be needed for some aspects of kinetochore regulation (Grallert et al., 2015; Porter et al., 2013). This general point is illustrated nicely by our experiments on the RVSF motif in Knl1. The N-terminal position of this SLiM is not important for SAC silencing in the absence of microtubules, but the strict conservation at this position throughout evolution indicates there is an essential requirement that still remains to be discovered (fig.3c,d and supp.fig.3). Interestingly, the microtubule binding site on Knl1 was recently shown to overlap the PP1-binding SLiMs (Bajaj et al., 2018). Therefore, we speculate that microtubules might need to compete with PP1 at the N-terminus. For example, they may need to elongate the Knl1 structure and disrupt PP1 binding at the same time. It will be interesting to test how this competition could impact on Aurora B regulation, error-correction and tension-sensing.

Although other properties of PP1 and PP2A could be important in some contexts, their inverse phospho-dependencies are clearly the defining features with regards to the SAC and kinetochore-microtubule attachments. This explains why the relevant phosphorylation sites are so well-conserved within each kinetochore SLiM (fig.3b and (Suijkerbuijk et al., 2012)). What then, is the relevance of the particular kinase inputs needed to phosphorylate these SLiMs? As discussed previously, Aurora B may regulate the PP1 SLiMs to allow the SAC phosphatase to respond to tension. We speculate that Plk1 and Cdk1 may similarly regulate the PP2A-B56 SLiM to allow the kinetochore-microtubule phosphatase to respond to microtubule attachment. This is predicted given that both kinases are recruited to kinetochores in an attachment-sensitive manner (Alfonso-Perez et al., 2019; Lenart et al., 2007; Liu et al., 2012). In fact, both are also recruited to the KMN network in a phosphorylation-dependent manner: Cyclin B/Cdk1 interacts with Mad1, a phospho-dependent interactor of Bub1, and Cdk1 can phosphorylate Bub1 to recruit Plk1 (Alfonso-Perez et al., 2019; Saurin, 2018). Interestingly, the key Mad1-Bub1 interaction has also been shown to be negatively regulated by kinetochore PP2A-B56 (Qian et al., 2017). Therefore, PP2A may also counteract Cdk1 and Plk1 activity to create yet more negative feedback to restrict its activity. This would serve to restrict PP2A from silence the SAC silencing even more, which could allow the current mathematical model to better approximate the real data. It should be stressed that this model is just a basic framework to explore how inverse phospho-dependencies affect phosphatase behaviour. In future, details of how the various kinase inputs respond to phosphatase activity should be layered into this model to provide a more complete picture of signal integration at the KMN network.

Finally, if kinetochores have selected to interact with PP1 and PP2A-B56 because of their inverse phospho-dependencies, then many other pathways have likely exploited the same features. This would explain the prevalence of phosphorylation sites within validated and predicted RVxF and LxxIxE SLiMs (fig.6c-e and supp.table.1). The invariant SLiM residues place some constraints on the type of kinase inputs that are tolerated within the motifs themselves (supp.fig.7), but phosphorylation outside of these regions can also regulate phosphatase binding (Kumar et al., 2016; Qian et al., 2015). Furthermore, PP1 uses co-operative interaction with other SLiMs, and some of these, such as the SILK motif in Knl1, are also phospho-inhibitable (Liu et al., 2010). Therefore, these additional interactions could allow a wide range of kinase inputs to converge and fine-tune PP1 binding strength. It will be important in future to fully characterise all possible SLiM-interactors for both PP1 and PP2A-B56, and to determine how these can be regulated by different kinases. As pointed out recently, kinases and phosphatases work together in many different ways to generate the right type of signal response (Gelens et al., 2018; Gelens and Saurin, 2018). Therefore, the ability of different phosphatases to positively or negatively couple to phosphorylation inputs represents a fundamental, but still poorly understood, aspect of signal integration.

## Supporting information

Supplementary Modelling Methods

## Acknowledgements

MHC, RJS and ATS are funded by Cancer Research UK (C47320/A21229 and C10988/A22566 to ATS). GV was funded by Tenovus Scotland (T14-19 to ATS). AC is funded by a grant of the Italian Association for Cancer Research (IG 21556). NED was supported by an SFI Starting Investigator Research Grant [13/SIRG/2193]. We thank staff at the Dundee Imaging Facility, the Flow Cytometry and Cell Sorting facility, and the Genetic Core Services Unit. We also thank Stephen Taylor for providing the HeLa Flp-in cell line and Geert Kops and Iain Cheeseman for antibodies.

## Author contributions

ATS, RJS and MHC conceived the study, designed the experiments and interpreted the data. RJS and MHC performed the majority of these experiments, with contributions from GV. FG and AC performed the modelling for figure 5. ND performed SLiM analysis for figure 6 and supplementary table 1. ATS wrote the manuscript with input from all authors.

## Declaration of interests

The authors declare no competing interests.

## Materials and Methods

### Cell culture and reagents

All cell lines were derived from HeLa Flp-in cells (a gift from S Taylor, University of Manchester, UK) (Tighe et al., 2008), which were authenticated by STR profiling (Eurofins). The cells were cultured in DMEM supplemented with 9% FBS and 50 µg/ml penicillin/streptomycin. During fluorescence time-lapse analysis, cells were cultured in Leibovitz’s L-15 media (900 mg/L D+ Galactose, 5mM Sodium Pyruvate, no phenol red) or DMEM (no phenol red) supplemented with 9% FBS and 50µg/ml penicillin/streptomycin. Doxycycline (1µg/ml) and thymidine (2 mM) was purchased from Sigma-Aldrich, nocodazole (3.3 µM) from Millipore, puromycin and hygromycin B from Santa Cruz biotechnology, MG132 (10 µM) from Selleckbio, AZ-3146 from Axon, ZM-447439 from Cayman Chemicals, RO-3306 from Tocris. Cells were screened every 4-8 weeks to ensure they were mycoplasma free.

### Plasmids and cloning

pcDNA5-YFP-BubR1^WT^ expressing an N-terminally YFP-tagged and siRNA-resistant wild-type BubR1 and pcDNA5-YFP-BubR1^ΔPP2A^ (also called BubR1^ΔKARD^), lacking amino acids 663-680 were described previously (Nijenhuis et al., 2014). All the remaining YFP-BubR1 mutants were subcloned by PCR amplification of DNA fragments followed by a Gibson assembly reaction to insert back into the original vector (pcDNA5-YFP-BubR1^WT^), except when indicated. pcDNA5-YFP-BUBR1 ^ΔCT^, was subcloned from pcDNA5-YFP-BubR1^WT^ by PCR amplification of the BubR1 fragment (excluding amino acids 664-1050). pcDNA5-YFP-BubR1^Long^ was constructed by insertion of a flexible 36 amino acid [GSG]-linker between amino acids 663 and 664. pcDNA5-YFP-BubR1^deP-PP2A^ was generated by site directed mutagenesis of the pcDNA5-YFP-BubR1^WT^ vector mutating the KARD motif from 5’-SIKKLSPIIEDSR-3’ to 5’-RSSTLPTIHEEEE-3’ (Kruse et al., 2018).

pcDNA5-YFP-Knl1^WT^ expressing an N-terminally YFP-tagged and siRNA-resistant wild-type Knl1, pcDNA5-YFP-Knl1^PP1(2A)^ (with S25A and S60A mutations, also called Knl1^2SA^) and pcDNA5-YFP-Knl1^ΔPP1^ (with RVSF at amino acids 58-61 mutated to AAAA, also called Knl1^4A^) were described previously (Nijenhuis et al., 2014). Site directed mutagenesis was used to further improve the resistance of pcDNA5-YFP-Knl1^WT^ construct to dsiRNA by modifying two extra codons (new dsiRNA-resistant site 5’-GCACGTGAGCTTGAAGGAA-3’, nucleotides 2678-2676). While confirming the accuracy of the mutagenesis by sequencing, we detected a deletion of amino acids 910 to 1120 in the Knl1 constructs used previously (Nijenhuis et al., 2014). This occurred between identical MELT13/17 sequences and was probably caused by recombination of the plasmid during bacterial culture. To correct this, a 3991 bp fragment (corresponding to nucleotides 2730-6720 in Knl1^WT^ plasmid) was amplified from genomic DNA of RPE cells and subcloned into the pcDNA5-YFP-Knl1^WT^ plasmid already containing the improved siRNA-resistant site. Gibson assembly was then performed to correct all Knl1 constructs by replacing the N-terminus through XhoI and PmlI restriction sites using pcDNA5-YFP-Knl1^PP1(2A)^ and pcDNA5-YFP-Knl1^ΔPP1^ as PCR templates. pcDNA5-YFP-Knl1 ^ΔNT^ (with deletion of the first 70 amino acids of Knl1) was also created by Gibson assembly using the same restriction sites.

pcDNA5-YFP-B56_γ1_-Knl1^ΔNT^ (Knl1 ^ΔNT^ fused to B56γ_1_) and pcDNA5-YFP-B56 _γ1_^CD^-Knl1^ΔNT^ (Knl1^ΔNT^ fused to a version of B56_γ1_ with a S296D mutation that disrupts PP2A binding (Vallardi et al., 2018)) were produced by restriction cloning using fragments generated by PCR from pcDNA5-YFP-B56_γ1_ and pcDNA5-YFP-B56_γ1_^CD^, respectively (Vallardi et al., 2018), both were inserted using NotI and KasI restriction sites creating a 28 amino acid linker before Knl1 ^ΔNT^. To create pcDNA5-YFP-Knl1 ^ΔNT-PP1mid^ a fragment containing the PP1-binding SILK and RVSF motifs (amino acids 24 to 70 of Knl1^WT^) was inserted at a BlpI site of pcDNA5-YFP-Knl1 ^ΔNT^ by Gibson assembly (between MELT-10 and MELT-11). pcDNA5-YFP-BubR1^ΔCT-PP1:Knl1^ was created by Gibson assembly of two PCR fragments: pcDNA5-YFP-BubR1^1-663^ (amplified from pcDNA5-YFP-BUBR1^WT^) and a fragment containing the first 70 amino acids of the N-terminal tail of Knl1 with the Aurora B sites mutated in the SLiM (amplified from pcDNA5-YFP-Knl1^PP1(2A)^) (a 6 amino acid linker connects BubR1^ΔCT^ to the N-terminal tail of Knl1). The same subcloning strategy was used to create pcDNA5-YFP-BubR1^ΔCT-Knl1^ but using a mutated N-terminal tail of Knl1 which cannot recruit PP1 (amplified from pcDNA5-YFP-Knl1^ΔPP1^). pcDNA5-YFP-BubR1^ΔCT^-B56_γ1_ was also subcloned with the same strategy but using a fragment containing B56 amplified from pcDNA5-YFP-B56_γ1_ (Vallardi et al., 2018), inserting B56_γ1_ and a 7 amino acid linker after amino acid 658 of BubR1.

pcDNA4-mTurquoise2(Turq2)-BubR1^WT^ was created by Gibson assembly of 3 PCR fragments: pcDNA4 backbone, a fragment containing BubR1^WT^ (amplified from pcDNA5-YFP-BubR1^WT^) and a fragment containing mTurquoise2. pcDNA5-Turq2-BubR1 ^deP-PP2A^ was created by restriction cloning using Acc65I and BstBI to replace the YFP originally present in pcDNA5-YFP-BubR1 ^deP-PP2A^ (Turq2 subcloned from pcDNA4-Turq2-BubR1^WT^). Similarly, Turq2-tagged version of BubR1^ΔCT^-B56_γ1_ was created by restriction cloning using NheI and NotI to remove YFP. A pcDNA4-Turq2 version of BubR1^ΔCT^ was created by restriction cloning using BstBI and HpaI to replace BubR1^WT^ from the original pcDNA4-Turq2-BubR1^WT^ vector. In the same way, pcDNA4-Turq2-BubR1^Long^ was created by restriction cloning using BstBI and Bsu36I, and pcDNA4-Turq2 versions of BubR1^ΔCT-PP1:Knl1^ and BubR1^ΔCT-Knl1^ were created by restriction cloning using BstBI and NotI. pMESV_Ψ_-mCherry-B56_γ1_ was produced by Gibson assembly of pMESV_Ψ_-mCherry backbone (amplified from pMESV_Ψ_-mCherry-CenpB-Mad1 (Maldonado and Kapoor, 2011)) and B56_γ1_ (amplified from pcDNA5-YFP-B56_γ1_) (Vallardi et al., 2018). All plasmids were fully sequenced to verify the transgene was correct.

### Gene expression

HeLa Flp-in cells stably expressing doxycycline-inducible constructs were derived from the HeLa Flp-in cell line by transfection with the relevant pcDNA5/FRT/TO vector and the Flp recombinase pOG44 (Thermo Fischer). Cells were subsequently selected in media containing 200 µg/ml hygromycin B for at least 2 weeks to select for stable integrants at the FRT locus. Cells expressing mCherry-B56γ1 in combination with YFP-tagged Knl1^WT/ΔPP1^ were generated by viral-integration of pMESV_Ψ_-mCherry-B56_γ1_ construct into the genome of HeLa Flp-in cells, followed by puromycin selection (1µg/ml) and were then transfected as above with YFP-Knl1^WT/ΔPP1^. Double mutant analysis was performed using cells that express a combination of Turq2-tagged BubR1 and YFP-tagged Knl1. These were generated by transient transfection of Turq2-tagged constructs into cells that were stably expressing doxycycline-inducible YFP-tagged recombinant proteins (generated as described above). These Turq2-tagged constructs were transfected 32 hours prior to endogenous gene knock-down (described below) and at least 72 hours prior to imaging or fixation. Plasmids were transfected into HeLa Flp-in cells using Fugene HD (Promega) according to the manufacturer’s instructions.

### Gene knockdown

For all experiments involving recombinant BubR1 or Knl1 in HeLa Flp-in cells, the endogenous mRNA was knocked down and replaced with an siRNA-resistant mutant. To knockdown endogenous BubR1 or Knl1 or both together, cells were transfected with 20nM BubR1 siRNA (5′-AGAUCCUGGCUAACUGUUC-3’) (Sigma-Aldrich) or 20nM Knl1 dsiRNA (5’-ATGCATGTATCTCTTAAGGA AGATGAA-3’) (Integrated DNA technologies) or both simultaneously for 16 h after which the cells were arrested in early S phase by addition of thymidine for 24 h. All siRNAs were transfected using Lipofectamine^®^ RNAiMAX Transfection Reagent (Life Technologies) according to the manufacturer’s instructions. BubR1 and Knl1 construct expression was induced by the addition of doxycycline during and following the thymidine block. After thymidine block, cells were release into media supplemented with doxycycline and, where appropriate, nocodazole for 5-7 hours for live imaging or 8.5 hours for fixed analysis. Mps1 and Aurora B were inhibited by adding AZ-3146 and/or ZM-447439 shortly prior to live cell imaging. For kinase inhibition in cells analysed by immunofluorescence, we nocodazole and MG132 was added first for 30 minutes, followed by a time-course of AZ-3146 and/or ZM-447439 with nocodazole and MG132.

### Immunofluorescence

Cells, plated on High Precision 1.5H 12-mm coverslips (Marienfeld), were fixed with 4% paraformaldehyde (PFA) in PBS for 10 min or pre-extracted (when using pRVSF-Knl1 or mCherry antibodies) with 0.1% Triton X-100 in PEM (100 mM Pipes, pH 6.8, 1 mM MgCl2 and 5 mM EGTA) for 1 minute before addition of 4% PFA for 10 minutes. Coverslips were washed with PBS and blocked with 3% BSA in PBS + 0.5% Triton X-100 for 30 min, incubated with primary antibodies overnight at 4°C, washed with PBS and incubated with secondary antibodies plus DAPI (4,6-diamidino2-phenylindole, Thermo Fischer) for an additional 2-4 hours at room temperature in the dark. Coverslips were then washed with PBS and mounted on glass slides using ProLong antifade reagent (Molecular Probes). All images were acquired on a DeltaVision Core or Elite system equipped with a heated 37°C chamber, with a 100×/1.40 NA U Plan S Apochromat objective using softWoRx software (Applied precision). Images were acquired at 1×1 binning using a CoolSNAP HQ or HQ2 camera (Photometrics) and processed using softWorx software and ImageJ (National Institutes of Health). All immunofluorescence images displayed are maximum intensity projections of deconvolved stacks and were chosen to most closely represent the mean quantified data.

### Image quantification

For quantification of immunostainings, all images of similarly stained experiments were acquired with identical illumination settings and cells expressing comparable levels of exogenous protein were selected and analysed using ImageJ. An ImageJ macro was used to threshold and select all kinetochores and all chromosome areas (excluding kinetochores) using the DAPI and anti-kinetochore antibody channels, as previously (Saurin et al., 2011). This was used to calculate the relative mean kinetochore intensity of various proteins ([kinetochores-chromosome arm intensity (test protein)] / [kinetochores-chromosome arm intensity (CENP-C)].

### Chromosome alignment assays

To observe chromosome alignment, cells were released from thymidine block for 7 hours before being synchronised at the G2/M boundary with a 2-hour treatment with RO-3306 (10μM). Cells were then washed three times and incubated for 15 minutes with full growth media before addition of MG132 for 30 minutes to prevent mitotic exit. Cells were then fixed and stained as described above and scored based on the number of misaligned chromosomes as aligned (0 misaligned chromosomes), mild (1-2), moderate (3-5) or severe (>6). This protocol is important because mutants that cause a prolonged arrest can otherwise cause cohesion fatigue, which skews the alignment data.

### Time-lapse analyses

For fluorescence imaging, cells were imaged in 8-well chamber slides (ibidi) in Leibovitz L-15 media with a heated 37°C chamber or in DMEM (no phenol red) with a heated 37°C chamber in 5% CO_2_. Images were taken every 4 minutes with either a 20× /0.4 NA air objective using a Zeiss Axio Observer 7 with a CMOS Orca flash 4.0 camera at 4×4 binning or a 40×/1.3 NA oil objective using a DV Elite system equipped with Photometrics CascadeII:1024 EMCCD camera at 4×4 binning. For brightfield imaging, cells were imaged in a 24-well plate in DMEM in a heated chamber (37°C and 5% CO_2_) with a 10×/0.5 NA objective using a Hamamatsu ORCA-ER camera at 2×2 binning on a Zeiss Axiovert 200M, controlled by Micro-manager software (open source: https://miro-manager.org/) or with a 20×/0.4 NA air objective using a Zeiss Axio Observer 7 as detailed above. Mitotic exit was defined by cells flattening down in the presence of nocodazole and Mps1 inhibitor. In assays where both recombinant BubR1 and Knl1 are expressed in cells, cells were selected for quantification based on high levels of Turq2 as an indication of successful transient transfection of Turq2-BubR1 constructs into cells stably expressing YFP-tagged Knl1 constructs.

### Antibodies

All antibodies were diluted in 3% BSA in PBS. The following primary antibodies were used for immunofluorescence imaging (at the final concentration indicated): chicken α-GFP (ab13970 from Abcam, 1:5000), mouse α-GFP (clone 4E12/8, a gift from P. Parker, 1:1000), rabbit α-pNDC80 Serine 55 (1:1000, GeneTex), guinea pig α-Cenp-C (BT20278 from Caltag + Medsystems), rabbit α -Bub1 (A300-373A from Bethyl, 1:1000), rabbit α -BubR1 (A300-386A from Bethyl, 1:1000), rabbit α-mCherry (GTX128508, Genetex,1:1000). The rabbit α-pMELT-Knl1 antibody is directed against Thr 943 and Thr 1155 of human Knl1 (Nijenhuis et al., 2014) (Gift from G.Kops, Hubrecht, NL). The pRVSF-Knl1 (pSer60-Knl1) antibody (custom rabbit polyclonals, gift from I. Cheeseman, MIT, USA) was used at 1:2000 dilution (Nijenhuis et al., 2014). Secondary antibodies used were highly-cross absorbed goat, α-chicken, α-rabbit, α-mouse or a-guinea pig coupled to Alexa Fluor 488, Alexa Fluor 568, or Alexa Fluor 647 (Thermo Fischer).

### Mathematical Modelling

The model consists of a set of ordinary differential equations (ODEs) that correspond to the diagram in Figure 5a. All binding/dissociation and phosphorylation/dephosphorylation reactions were modelled according to simple mass-action kinetics. The assumption of no substrate specificity means that PP1 and PP2A act on the same substrates (pMELT, pRVSF, pBubR1, and pNdc80) and that for each substrate there is only one parameter describing the catalytic activities of both. We assume that the total amounts for all species are conserved, except for Mps1 and Aurora B whose amounts can be changed externally to simulate the experimental inhibition of kinases. Parameters were determined by fitting the model to the data from Fig. 1c-f and Supp. Fig. 6. For detailed information see Supplementary Methods.

### Motif analysis

A set of PP1-binding peptides for the RVxF-binding pocket and PP2A-B56-binding peptides for the LxxIxE-binding pocket were created from experimentally validated peptides in the PP1 and PP2A literature. The dataset contained 110 RVxF and 27 LxxIxE motifs. A position-specific scoring matrix (PSSM) was constructed from each set of peptides based on amino acid frequencies weighted using peptide similarity weights and pseudocounts using the PSI BLAST IC scoring scheme as defined in the PSSMSearch tool (Altschul et al., 1997; Krystkowiak et al., 2018). Each PSSM was screened against the human UniProt reviewed proteins using PSSMSearch (Krystkowiak et al., 2018) and filtered using PSSM score *p-value* with a cut-off of 0.0001, taxonomic range based on conservation of the consensus outside the mammalian clade, localisation based on intracellular localisation GO terms, and accessibility based on: (i) overlap with a resolved region in a structure from PDB, (ii) intrinsic disorder predictions (retaining only peptides found in disordered regions as defined by an IUPred score < 0.3 [15769473]) and (iii) UniProt annotation of topologically inaccessible regions (e.g. transmembrane and extracellular regions) [25348405]. Applying these criteria, we produced sets of predicted 344 RVxF-binding and 210 PP2A-B56-binding motifs (Supplementary Table 1). The phosphorylated (experimentally validated phosphorylation sites annotated in the UniProt, phospho.ELM or phosphosite databases) or phosphorylatable (any serine or threonine) residues within the predicted and validated sets were collected and the kinase specificity of each site was annotated as basophilic ([KR]xS or [KR]xxS), acidophilic ([DEN]x[ST]) or proline-directed ([ST]P). Enrichment of motif specificity determinants were calculated as the binomial probability (*prob*^*aa*^ = *binomial(k,n,p)*) where *k* is the observed residue count at each position for a residue, *n* is the number of the instances of motifs and p is the background amino acid frequency of a residue based on the disordered regions of the human proteome. Enrichment of groupings (KR - basic, DE - acidic, ST - phosphorylatable by serine/threonine kinases) was calculated similarly. RVxF-binding motifs are available at http://slim.ucd.ie/motifs/pp1/index.php?page=instances and PP2A-B56-binding motifs are available at http://slim.ucd.ie/pp2a/index.php?page=instances

### Statistics

Two-tailed, unpaired *t*-tests with Welch’s correction were performed to compare the means values between experimental groups in immunofluorescence quantifications (using Prism 6 software).

