## Supplementary Figures for "PP1 and PP2A use opposite phospho-dependencies to control distinct processes at the kinetochore"

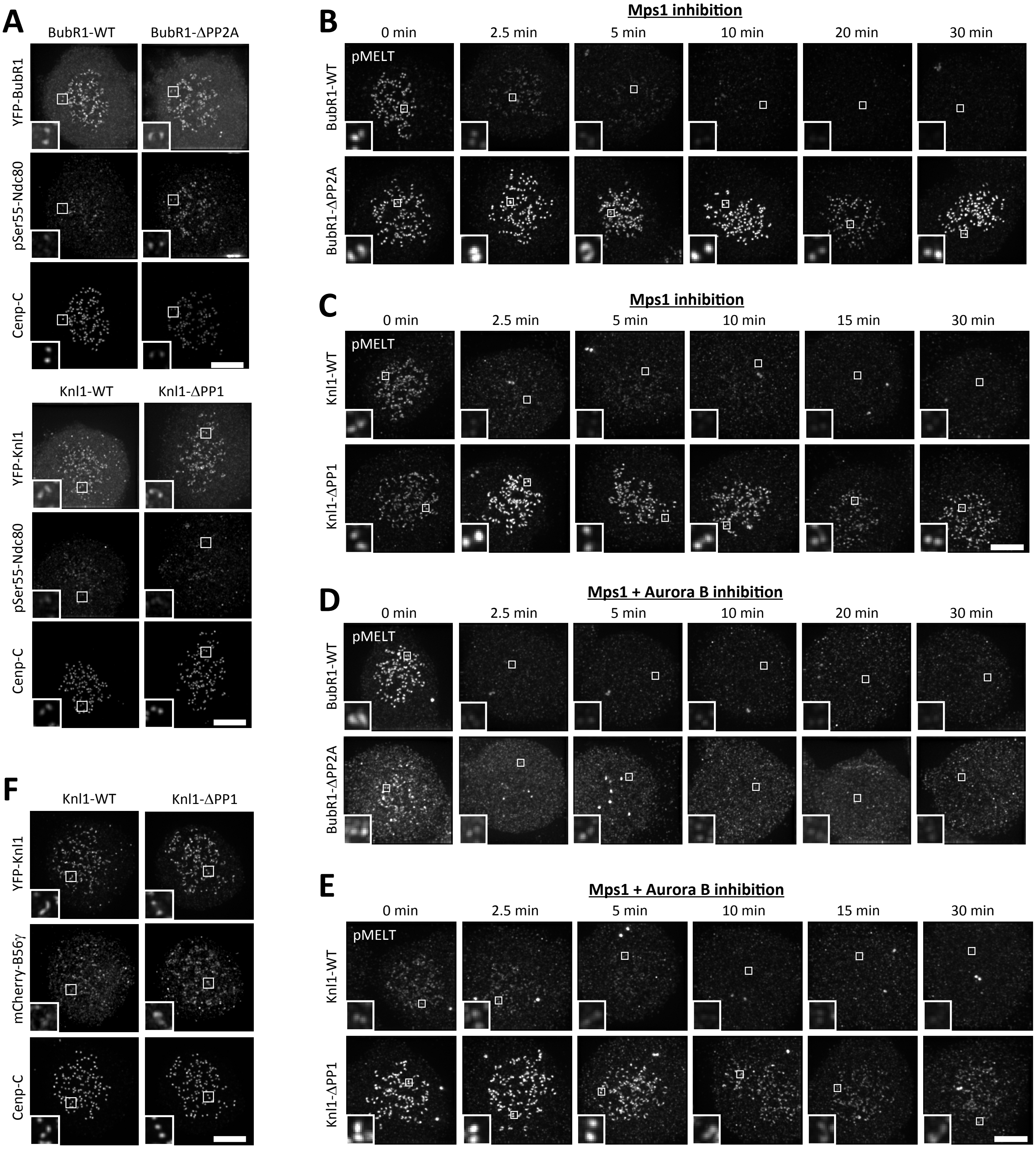


Supplementary Figure 1. Immunofluorescence images to show that PP1-Knl1 and PP2A-B56 exert control over different pathways and processes at the kinetochore.

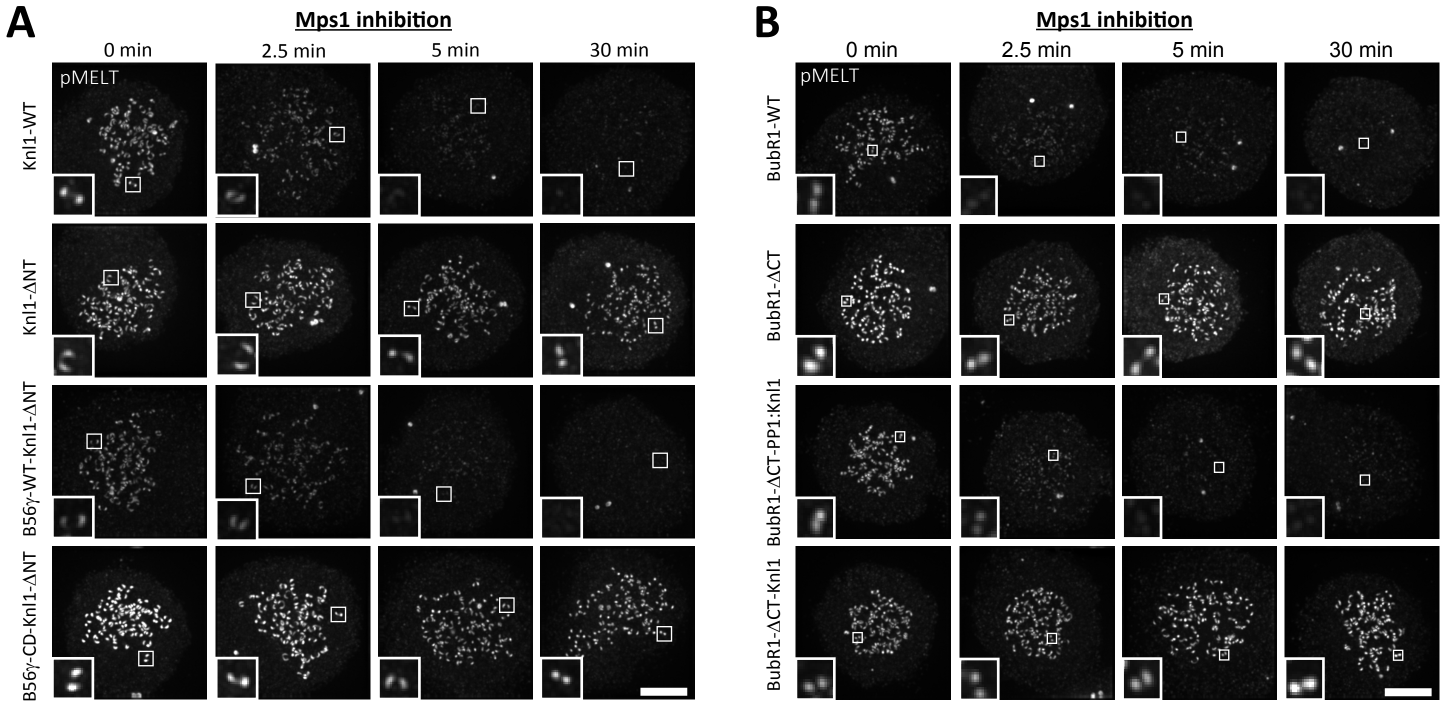


Supplementary Figure 2. Immunofluorescence images to show that PP1-Knl1 and PP2A-B56 can functionally substitute for each other at the kinetochore.

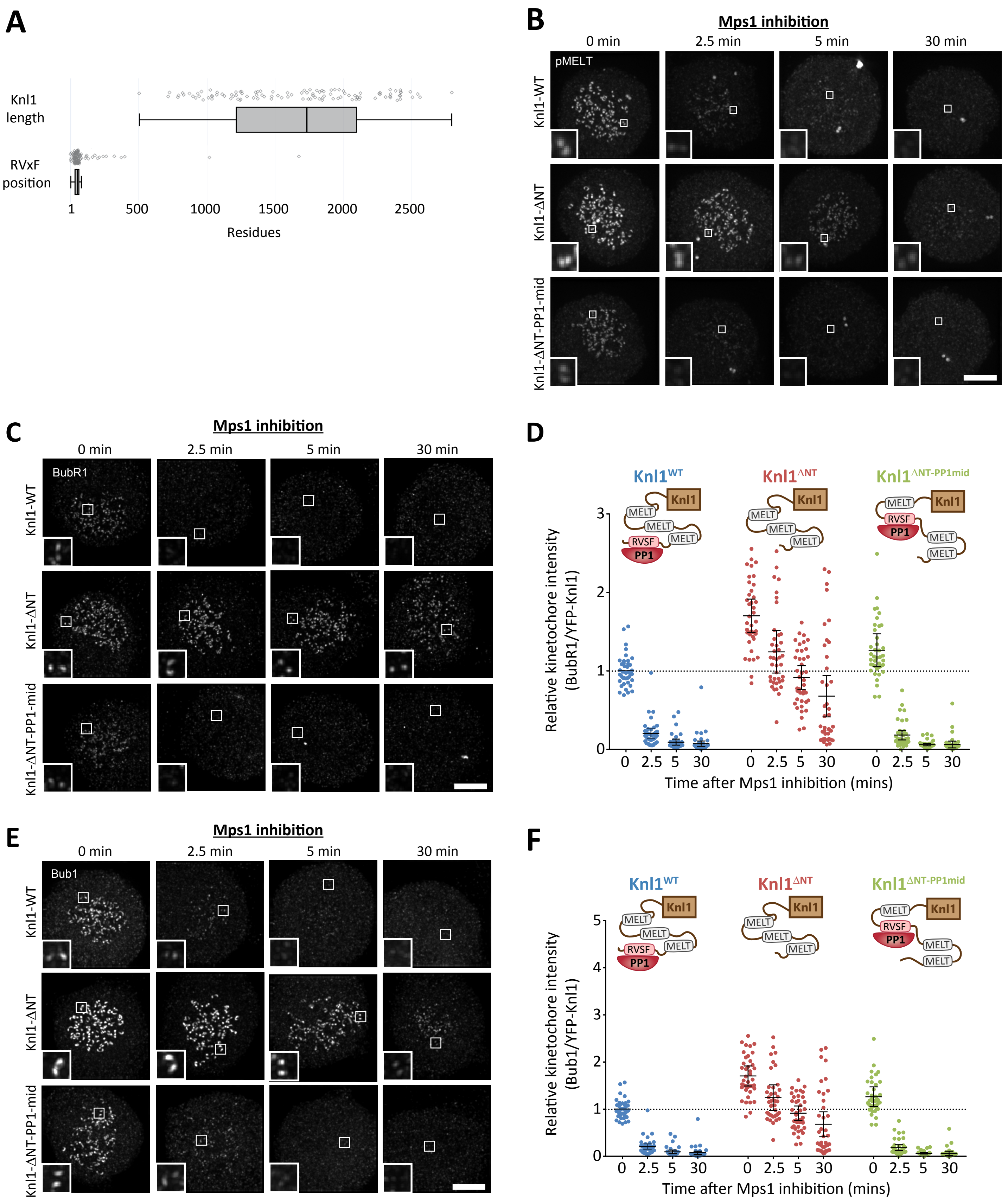


Supplementary Figure 3. The PP1 binding SLiM is conserved at the N-terminus of Knl1 and this position is not critical for MELT dephosphorylation in nocodazole.

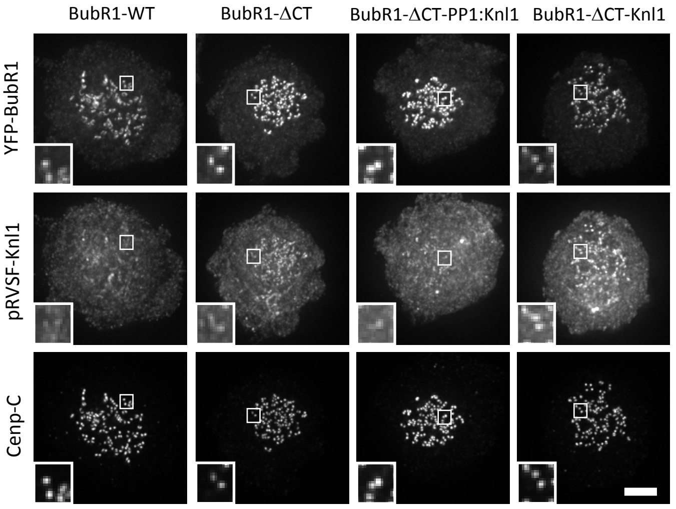


Supplementary Figure 4. Immunofluorescence images to show that PP1 can dephosphorylate KNL1-RVSF when recruited to BubR1.

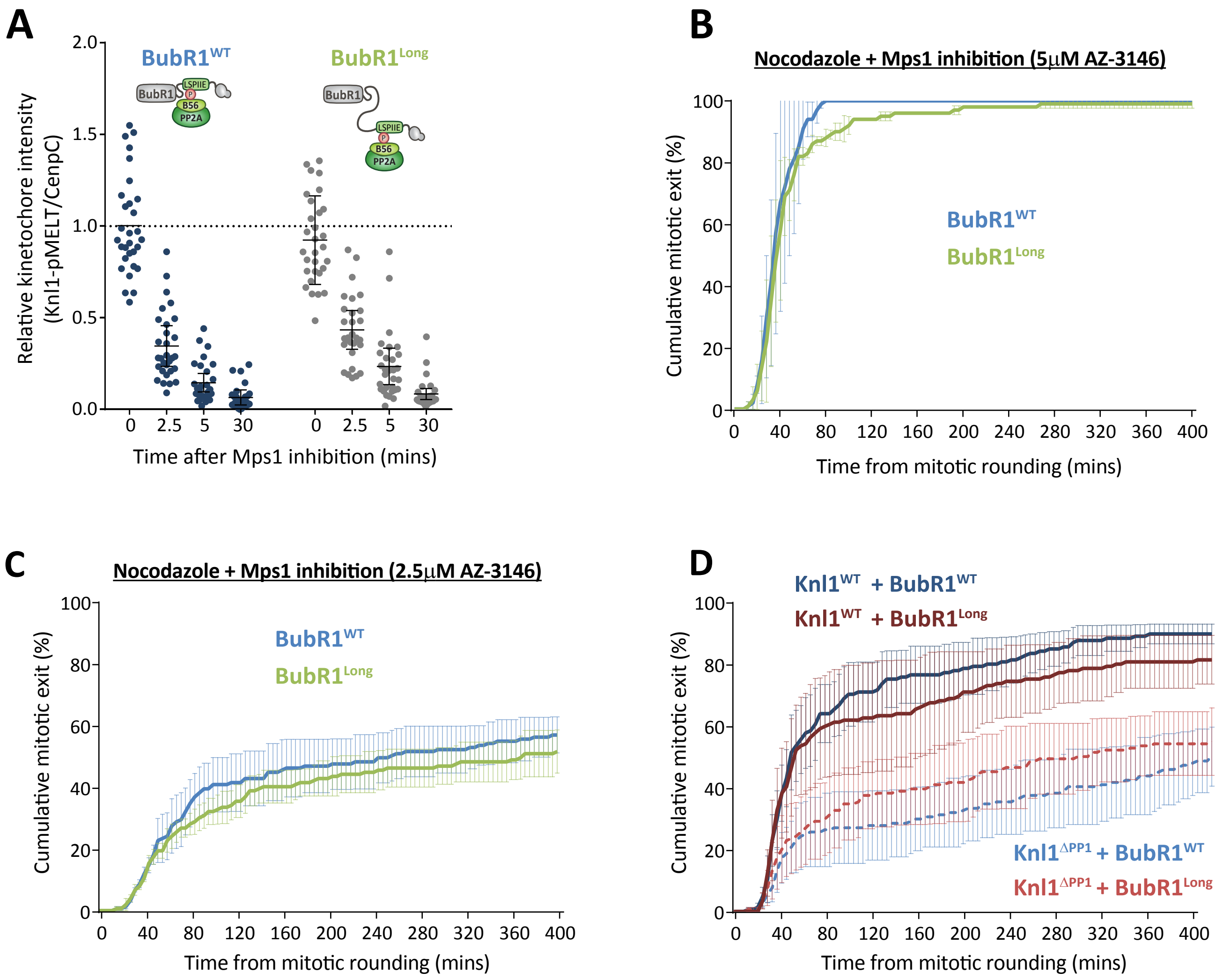


Supplementary Figure 5. Insertion of a long flexible linker before the LxxIxE motif in BubR1 does not improve the ability of PP2A to silence the SAC.

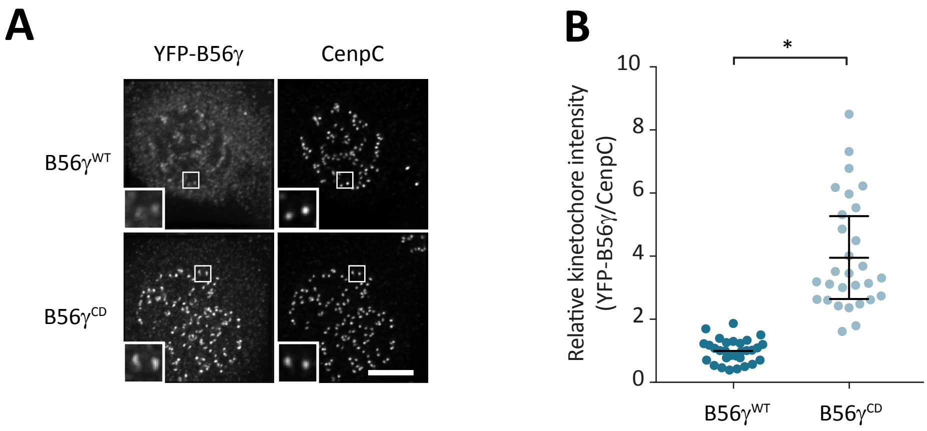


Supplementary Figure 6. Effect of negative feedback on B56γ kinetochore levels.

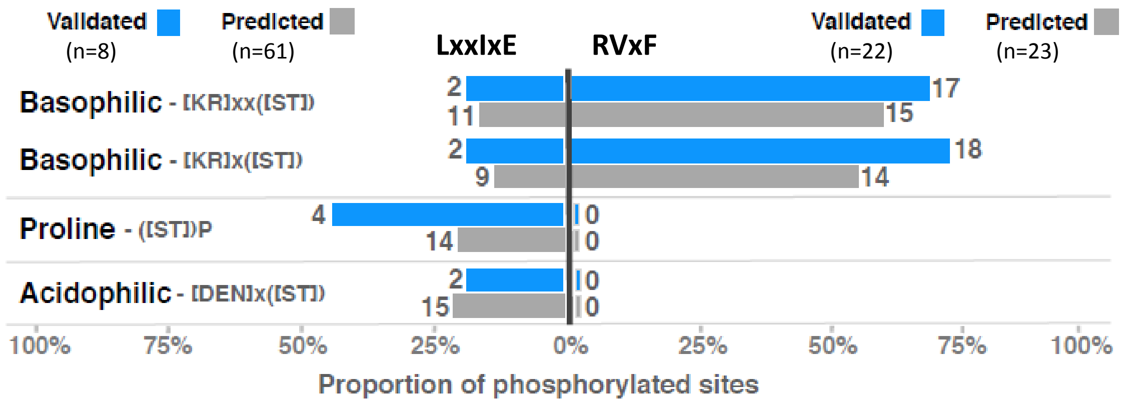


Supplementary figure 7. Summary of the kinase specificity determinants overlapping the phosphorylated and phosphorylatable sites in the experimentally validated LxxIxE and RVxF motifs.
