## Supplementary Modelling Methods for "PP1 and PP2A use opposite phospho-dependencies to control distinct processes at the kinetochore"

### Mathematical Model: Materials and Methods

February 27, 2019

#### 1 Modelling approach, terminology and assumptions

The model consists of a set of ordinary differential equations (ODEs) that correspond to the diagram in Fig. 5a. All binding/dissociation and phosphorylation/dephosphorylation reactions were modeled according to simple mass-action kinetics.

In the following, "[X]" stands for the amount of species X. If X is an 'atomic' species (i.e. corresponding to one of the individual rectangular boxes in Fig. 5a), then "$[X_{tot}]$" denotes the total amount of species X, including phosphorylated forms and all complexes that contain X as a component, while "$[X_{free}]$" stands for the amount of X that is unbound and unphosphorylated. "pX" stands for the phosphorylated form of X and "X : Y" for a form in which species X and Y are bound. Binding/dissociation rates for species X are referred to as "$kb_X$" and "$kd_X$" and phosphorylation/dephosphorylation rates as "$kp_X$" and "$kdp_X$", respectively.

Using ODEs means that the behaviour of individual molecules is not represented and that all species are assumed to be well-mixed. Thus, effectively the model describes the processes occuring on all Knl1 molecules located on one representative kinetochore. We assume that the total amounts for all species are conserved, except for Mps1 and AurB whose amounts can be changed externally to simulate the experimental inhibition of kinases. Furthermore, we assume that BubR1 can be phosphorylated both in its free form and when bound to pMELT but allow that this may happen with different rates (corresponding to the parameters $kp_{BubR1(free)}$ and $kp_{BubR1}$). PP1 and PP2A are assumed to be catalytically active only when bound to Knl1 (via RVSF and pMELT : pBubR1, respectively. In reality, PP2A can also bind to unphosphorylated BubR1, but we neglect this possibility because phosphorylation by Cdk1 and Plk1 increases the binding affinity by 11-fold and 5-fold, respectively [1]. To keep the model as simple as possible we assumed that one phosphorylation event (Cdk1) is needed to allow PP2A binding. For brevity, the active phosphatases are denoted by "$PP1_{act}$" and "$PP2_{act}$".

The assumption of no substrate specificity means that PP1 and PP2A act on the same substrates (pMELT, pRVSF, pMELT : pBubR1, and pNdc80) and that for each substrate there is only one parameter describing the catalytic activites of both.

#### 2 Equations

The model can be described using nine linearly independent equations:

$$\frac{d\,[PP1_{act}]}{dt} = kb_{PP1} \cdot [RVSF] \cdot [PP1_{free}] - kd_{PP1} \cdot [PP1_{act}]$$

$$\frac{d\,[PP2A_{act}]}{dt} = kb_{PP2A} \cdot [PP2A_{free}] \cdot [pMELT : pBubR1] + kb_{BubR1} \cdot [pBubR1 : PP2A_{free}] \cdot [pMELT] - kd_{PP2A} \cdot [PP2A_{act}]$$

$$
\begin{aligned}
& -\mathrm{kd}_{\mathrm{BubR1}} \cdot [\mathrm{PP2A_{act}}] \\
\frac{d\,[\mathrm{pBubR1_{free}}]}{dt} =\; & \mathrm{kd}_{\mathrm{BubR1}} \cdot [\mathrm{pMELT:pBubR1}] \\
& +\mathrm{kd}_{\mathrm{PP2A}} \cdot [\mathrm{pBubR1:PP2A_{free}}] \\
& +\mathrm{kp}_{\mathrm{BubR1(free)}} \cdot [\mathrm{BubR1_{free}}] \cdot [\mathrm{Cdk1}] \\
& -\mathrm{kb}_{\mathrm{BubR1}} \cdot [\mathrm{pBubR1_{free}}] \cdot [\mathrm{pMELT}] \\
& -\mathrm{kb}_{\mathrm{PP2A}} \cdot [\mathrm{pBubR1_{free}}] \cdot [\mathrm{PP2A_{free}}]
\end{aligned}
$$

$$
\begin{aligned}
\frac{d\,[\mathrm{pBubR1:PP2A_{free}}]}{dt} =\; & \mathrm{kb}_{\mathrm{PP2A}} \cdot [\mathrm{pBubR1_{free}}] \cdot [\mathrm{PP2A_{free}}] \\
& +\mathrm{kd}_{\mathrm{BubR1}} \cdot [\mathrm{PP2A_{act}}] \\
& -\mathrm{kd}_{\mathrm{PP2A}} \cdot [\mathrm{pBubR1:PP2A_{free}}] \\
& -\mathrm{kb}_{\mathrm{BubR1}} \cdot [\mathrm{pBubR1:PP2A_{free}}] \cdot [\mathrm{pMELT}]
\end{aligned}
$$

$$
\begin{aligned}
\frac{d\,[\mathrm{pMELT}]}{dt} =\; & \mathrm{kd}_{\mathrm{BubR1}} \cdot [\mathrm{pMELT:BubR1}] \\
& +\mathrm{kd}_{\mathrm{BubR1}} \cdot [\mathrm{pMELT:pBubR1}] \\
& +\mathrm{kd}_{\mathrm{BubR1}} \cdot [\mathrm{PP2A_{act}}] \\
& +\mathrm{kp}_{\mathrm{MELT}} \cdot [\mathrm{MELT}] \cdot [\mathrm{Mps1}] \\
& -\mathrm{kb}_{\mathrm{BubR1}} \cdot [\mathrm{BubR1_{free}}] \cdot [\mathrm{pMELT}] \\
& -\mathrm{kb}_{\mathrm{BubR1}} \cdot [\mathrm{pBubR1_{free}}] \cdot [\mathrm{pMELT}] \\
& -\mathrm{kb}_{\mathrm{BubR1}} \cdot [\mathrm{pBubR1:PP2A_{free}}] \cdot [\mathrm{pMELT}] \\
& -\mathrm{kdp}_{\mathrm{MELT}} \cdot [\mathrm{pMELT}] \cdot ([\mathrm{PP1_{act}}] + [\mathrm{PP2A_{act}}])
\end{aligned}
$$

$$
\begin{aligned}
\frac{d\,[\mathrm{pMELT:BubR1}]}{dt} =\; & \mathrm{kb}_{\mathrm{BubR1}} \cdot [\mathrm{BubR1_{free}}] \cdot [\mathrm{pMELT}] \\
& +\mathrm{kdp}_{\mathrm{BubR1}} \cdot [\mathrm{pMELT:pBubR1}] \cdot ([\mathrm{PP1_{act}}] + [\mathrm{PP2A_{act}}]) \\
& -\mathrm{kd}_{\mathrm{BubR1}} \cdot [\mathrm{pMELT:BubR1}] \\
& -\mathrm{kp}_{\mathrm{BubR1}} \cdot [\mathrm{pMELT:BubR1}] \cdot [\mathrm{Cdk1}]
\end{aligned}
$$

$$
\begin{aligned}
\frac{d\,[\mathrm{pMELT:pBubR1}]}{dt} =\; & \mathrm{kb}_{\mathrm{BubR1}} \cdot [\mathrm{pBubR1_{free}}] \cdot [\mathrm{pMELT}] \\
& +\mathrm{kp}_{\mathrm{BubR1}} \cdot [\mathrm{pMELT:BubR1}] \cdot [\mathrm{Cdk1}] \\
& +\mathrm{kd}_{\mathrm{PP2A}} \cdot [\mathrm{PP2A_{act}}] \\
& -\mathrm{kd}_{\mathrm{BubR1}} \cdot [\mathrm{pMELT:pBubR1}] \\
& -\mathrm{kdp}_{\mathrm{BubR1}} \cdot [\mathrm{pMELT:pBubR1}] \cdot ([\mathrm{PP1_{act}}] + [\mathrm{PP2A_{act}}]) \\
& -\mathrm{kb}_{\mathrm{PP2A}} \cdot [\mathrm{PP2A_{free}}] \cdot [\mathrm{pMELT:pBubR1}]
\end{aligned}
$$

$$
\begin{aligned}
\frac{d\,[\mathrm{pRVSF}]}{dt} =\; & \mathrm{kp}_{\mathrm{RVSF}} \cdot [\mathrm{RVSF}] \cdot [\mathrm{AurB}] \\
& -\mathrm{kdp}_{\mathrm{RVSF}} \cdot [\mathrm{pRVSF}] \cdot ([\mathrm{PP1_{act}}] + [\mathrm{PP2A_{act}}])
\end{aligned}
$$

$$
\begin{aligned}
\frac{d\,[\mathrm{pNdc80}]}{dt} =\; & \mathrm{kp}_{\mathrm{Ndc80}} \cdot [\mathrm{Ndc80}] \cdot [\mathrm{AurB}] \\
& -\mathrm{kdp}_{\mathrm{Ndc80}} \cdot [\mathrm{pNdc80}] \cdot ([\mathrm{PP1_{act}}] + [\mathrm{PP2A_{act}}])
\end{aligned}
$$

### 3 Free amounts

The free amounts can be determined by the following algebraic relations, given that total amounts are conserved:

$$
\begin{aligned}
[\mathrm{PP1_{free}}] &= [\mathrm{PP1_{tot}}] \\
&\quad - [\mathrm{PP1_{act}}]
\end{aligned}
$$

$$
\begin{aligned}
[\mathrm{PP2A_{free}}] &= [\mathrm{PP2A_{tot}}] \\
&\quad - [\mathrm{PP2A_{act}}] \\
&\quad - [\mathrm{pBubR1 : PP2A_{free}}] \\
&\quad - [\mathrm{BubR1 : PP2A_{free}}]
\end{aligned}
$$

$$
\begin{aligned}
[\mathrm{BubR1_{free}}] &= [\mathrm{BubR1_{tot}}] \\
&\quad - [\mathrm{pBubR1_{free}}] \\
&\quad - [\mathrm{pMELT : BubR1}] \\
&\quad - [\mathrm{pMELT : pBubR1}] \\
&\quad - [\mathrm{pBubR1 : PP2A_{free}}] \\
&\quad - [\mathrm{BubR1 : PP2A_{free}}] \\
&\quad - [\mathrm{PP2A_{act}}]
\end{aligned}
$$

$$
\begin{aligned}
[\mathrm{MELT_{free}}] &= [\mathrm{MELT_{tot}}] \\
&\quad - [\mathrm{pMELT}] \\
&\quad - [\mathrm{pMELT : BubR1}] \\
&\quad - [\mathrm{pMELT : pBubR1}] \\
&\quad - [\mathrm{PP2A_{act}}]
\end{aligned}
$$

$$
\begin{aligned}
[\mathrm{RVSF_{free}}] &= [\mathrm{RVSF_{tot}}] \\
&\quad - [\mathrm{pRVSF}] \\
&\quad - [\mathrm{PP1_{act}}]
\end{aligned}
$$

$$
\begin{aligned}
[\mathrm{Ndc80_{free}}] &= [\mathrm{pNdc80_{tot}}] \\
&\quad - [\mathrm{pNdc80}]
\end{aligned}
$$

In order to compare the model results to the experimental measurements, we defined the total amount of phosphorylated MELT as an auxiliary variable in the following way:

$$
\begin{aligned}
[\mathrm{pMELT_{tot}}] &= [\mathrm{pMELT}] \\
&\quad + [\mathrm{pMELT : BubR1}] \\
&\quad + [\mathrm{pMELT : pBubR1}] \\
&\quad + [\mathrm{PP2A_{act}}]
\end{aligned}
$$

### 4 Total amounts

The fixed total amounts were chosen according to the following assumptions: First, there are multiple MELT motifs, but only one RVSF site on each Knl1 molecule. Second, the species that bind to these motifs are available in sufficient amounts to potentially 'saturate' the motifs. Finally, total amounts for PP1 and PP2A are equal.

The choice of units is arbitrary because we only used relative amounts when comparing the model output to experimental data, and for clarity we leave out all units in the following. The values for the external kinases (Mps1, AurB, and Cdk1) can be chosen arbitrarily without loss of generality because their activities are separately determined by fitting the respective parameters.

Given these assumptions, we chose the following values:

| species | amount |
|---|---|
| [$PP1_{tot}$] | 10 |
| [$PP2A_{tot}$] | 10 |
| [$MELT_{tot}$] | 10 |
| [$BubR1_{tot}$] | 10 |
| [$RVSF_{tot}$] | 1 |
| [$Ndc80_{tot}$] | 1 |
| [Mps1] | 1 |
| [AurB] | 1 |
| [Cdk1] | 1 |

### 5 Implementation of mutants and parameter optimization

The remaining parameters were determined by fitting simulated time courses of $pMELT_{tot}$ to two pieces of data. The first were the experimental data from Fig. 1c-f. More specifically, we started the simulations from the steady state corresponding to a condition of metaphase arrest ([Mps1] = [AurB] = 1) and set either [Mps1] = 0 or [Mps1] = [AurB] = 0. The WT condition corresponds to the model as defined above, the ΔPP2A and ΔPP1 mutants were implemented by setting [$PP2A_{tot}$] and [$PP1_{tot}$] = 0, respectively. Initial amounts for [$pMELT_{tot}$] were fit to be ≈ 50% of [$MELT_{tot}$] in WT, ≈ 80% in ΔPP2A, and ≈ 100% in ΔPP1. To be able to investigate the contribution of negative feedback in a more fine-grained way, parameters were additionally fit to the data in Supp. Fig. 6. Here, we considered [$PP2A_{act}$] as a readout for B56$\gamma$. The B56$\gamma^{CD}$ mutant was implemented by replacing all occurences of the sum [$PP1_{act}$] + [$PP2A_{act}$] in the above equations by [$PP1_{act}$].

Based on these data, we obtained the following parameter set:

| parameter | value |
|---|---|
| $kb_{PP1}$ | 0.13 |
| $kd_{PP1}$ | 0.20 |
| $kb_{PP2A}$ | 0.32 |
| $kd_{PP2A}$ | 1.86 |
| $kb_{BubR1}$ | 0.18 |
| $kd_{BubR1}$ | 14.21 |
| $kp_{BubR1}$ | 0.076 |
| $kp_{BubR1(free)}$ | 0.0076 |
| $kdp_{BubR1}$ | 3.35 |
| $kp_{MELT}$ | 0.37 |
| $kdp_{MELT}$ | 0.70 |
| $kp_{RVSF}$ | 72.91 |
| $kdp_{RVSF}$ | 11.95 |

For Fig. 5e the BubR1 – PP2A mutant was implemented by assuming that BubR1 phosphorylation and PP2A binding are quasi-instantaneous and irreversible (setting $kp_{BubR1} = kp_{BubR1(free)} = kb_{PP2A} = 10.000$ and $kdp_{BubR1} = kd_{PP2A} = 0$). The Knl1-PP2A mutant was implemented by setting $kb_{PP2A} = kd_{PP2A} = 0$ and using a fixed amount of [$PP2A_{act}$] = 1, which corresponds to one copy of PP2A per molecule of Knl1. The curves in Fig. 5e were generated by calculating the steady state value of [$\text{pMELT}_{\text{tot}}$] for different levels of [Mps1]. Sensitivity is commonly defined as the slope in a logarithmic plot [2].

### 6 Computational methods

The model was implemented in the Systems Biology Markup Language (SBML) and is available as a supplementary file. Parameter fitting and simulations were performed using the Python package SloppyCell [3, 4] and further analyzed using custom scripts written in Python.
